# A proteome-wide biochemical screen defines binding determinants of the core autophagy protein LC3B

**DOI:** 10.64898/2026.01.05.697675

**Authors:** Jennifer Kosmatka, Cong Liu, Jackson C. Halpin, Daniel Lim, Joseph H. Davis, Amy E. Keating

**Affiliations:** Department of Biology, Massachusetts Institute of Technology, Cambridge, Massachusetts, 02139, United States; Broad Institute of MIT and Harvard, Massachusetts Institute of Technology, Cambridge, Massachusetts, 02139, United States; Computational and Systems Biology Graduate Program, Massachusetts Institute of Technology, Cambridge, Massachusetts, 02139, United States; Department of Biological Engineering Massachusetts Institute of Technology, Cambridge, Massachusetts, 02139, United States

## Abstract

Human MAP1LC3B (LC3B) binds proteins involved in autophagy and other cellular processes using a degenerate four-residue short linear motif known as the LC3-interacting region (LIR). Biochemical and structural studies have identified LIRs in many LC3B interaction partners, but the sequence features that contribute to binding have not been systematically explored. To discover peptides that interact with LC3B and deeply profile the key binding determinants, we screened a library of ∼500,000 36-residue peptides derived from the human proteome using bacterial cell-surface display. Analysis of the screening data, coupled with structural studies and site-directed mutagenesis, revealed exceptions to the canonical LIR motif ([FWY]_0_-X_1_-X_2_-[LVI]_3_) and a strong preference for negatively charged residues adjacent to the LIR, which we visualized in a newly determined structure of LC3B bound to a peptide isolated in our screen. Guided by our screening data, we designed synthetic LIR-containing peptides that bind LC3B with affinities comparable to the tightest measured natural binder. Finally, we determined that mutations in LC3B commonly thought to abrogate binding of LIR-containing peptides instead alter LC3B binding specificity, leading to enhanced binding of some LIR-containing sequences. Taken together, our results refine the LIR motif definition, expand the network of candidate LC3B interaction partners, and highlight how mutations at the LIR docking site can modulate affinity and specificity.

## INTRODUCTION

Intrinsically disordered regions (IDRs) in proteins participate in key cellular processes via interactions with globular protein domains (Dyson and Wright 2005). <u>S</u>hort <u>li</u>near <u>m</u>otifs (SLiMs) mediate these interactions using contiguous stretches of 3 – 10 amino acids that undergo disorder-to-order transitions upon binding, typically resulting in interactions with equilibrium dissociation constants (K_D_) in the micromolar range (Tompa *et al*. 2014; Wright and Dyson 2015; Diella 2008; Davey *et al*. 2012; Van Roey *et al*. 2014). For most SLiMs, the sequence features necessary and sufficient for binding are incompletely understood. Cataloged regular expressions, such as those in the Eukaryotic Linear Motif (Kumar *et al*. 2022) and Motif Map of the Proteome (https://slim.icr.ac.uk/momap/) databases, capture the features that best typify known binders but also highlight the limitations of existing motif specifications, which are satisfied by many sequences that fail to bind (Bugge *et al*. 2020). Although the number of experimentally defined SLiMs has expanded in recent years through the targeted assessment of individual binding partners, more comprehensive, high-throughput approaches are required to refine these recognition patterns and to support the development of prediction tools (Davey *et al*. 2023; Kumar *et al*. 2024).

SLiMs mediate interactions in the conserved proteostasis pathway macroautophagy (Popelka 2020), hereafter referred to as autophagy. This pathway involves the *de novo* formation of a double-membrane organelle known as the autophagosome, which transports cellular contents to the lysosome for degradation (Mizushima *et al*. 2008). Microtubule-associated protein 1A/1B light chain 3B (LC3B) and its homologs are critical at multiple stages of the autophagic process (Rogov *et al*. 2023) and participate via interaction with many partners that contain a SLiM termed the LC3-interaction region (LIR) (Birgisdottir *et al*. 2013). During autophagy, hundreds of LC3B proteins are covalently linked to the growing autophagosome through reversible conjugation to phosphatidylethanolamine embedded in the autophagosomal membrane (Kabeya *et al*. 2004). LC3B•LIR complexes, enhanced by multivalency and avidity at the membrane surface (Sawa-Makarska *et al*. 2014; Wurzer *et al*. 2015; Lee and Davis 2024), tether cellular components to the autophagosome either directly as cargo or indirectly using selective autophagy receptors to ensure efficient degradation (Gubas and Dikic 2022). LC3B•LIR interactions are also critical for autophagosome biogenesis, trafficking, and lysosomal fusion (Johansen and Lamark 2020). Mutations that alter LC3B•LIR interactions have been linked to the development of many diseases, including neurodegenerative disorders, aging, and cancer (Ramesh Babu *et al*. 2008; Park *et al*. 2020; Fas *et al*. 2021; Brennan *et al*. 2022). LC3B also has important roles in LC3-associated forms of phagocytosis (Florey and Overholtzer 2012), endocytosis (Heckmann *et al*. 2019), and macropinocytosis (Sønder *et al*. 2021).

LC3B is one of six hAtg8 proteins that are homologous to the yeast Atg8 protein and that share a common architecture consisting of a ubiquitin-like β-grasp fold with two additional N-terminal α-helices (Sugawara *et al*. 2004). The 14 kDa LC3B and its paralogs bind to the information-poor consensus LIR [FWY]_0_-X_1_-X_2_-[LVI]_3_ (Chatzichristofi *et al*. 2023), where X can be any amino acid, via two conserved hydrophobic pockets that form the <u>L</u>IR-<u>d</u>ocking <u>s</u>ite (LDS). In its canonical binding mode, the aromatic residue [FWY]_0_ engages the first hydrophobic pocket (HP1), and the aliphatic residue [LVI]_3_ engages the second hydrophobic pocket (HP2) (Rogov *et al*. 2023). This degenerate LIR motif occurs ∼170,000 times in the human proteome, with ∼19,000 occurrences falling in regions of predicted disorder. It is unlikely that all of these sequences represent functional interaction sites; indeed, several proteins containing canonical LIRs fail to bind LC3B with measurable affinity (Chatzichristofi *et al*. 2023). Moreover, recent discoveries of binders that deviate from the canonical LIR motif have expanded the repertoire of hAtg8 binding modes, and thus increased the number of potential interactors. For example, Li *et al*. reported that LC3B binds to ankyrin-G using the LIR sequence ^1991^WIEF^1994^, wherein an aromatic Phe residue engages HP2 (Li *et al*. 2018), and Knævelsrud *et al*. found that sorting nexin-18 (SNX18) binds hAtg8 paralogs using an expanded five-residue motif (154WDDEW158) (Knævelsrud *et al*. 2013). The proteome-wide prevalence of binders that lack the canonical [FWY]_0_-X_1_-X_2_-[LVI]_3_ motif remains unknown.

Given that not all instances of the LIR motif bind to LC3B, and that not all LC3B interactors contain a LIR motif, we sought to understand determinants beyond the core LIR that contribute to binding. To discover these sequence determinants, we used a bacterial display assay to screen ∼500,000 peptides derived from the human proteome for those capable of binding to LC3B. We determined the binding affinity of 45 peptides from the screen and elucidated binding mechanisms that support SLiM-LC3B interactions for known and novel binders through structural and biochemical analysis of identified peptides and mutated variants. Our results expand the number and types of residues that can engage HP2 on LC3B and strongly support the role of N-terminal acidic residues in enhancing LIR•LC3B binding affinity, allowing us to design a synthetic LIR peptide with affinity comparable to that of the tightest known native LC3B binder.

## RESULTS

### A high-throughput bacterial display screen identifies thousands of LC3B-binding peptides

To rapidly identify peptides capable of binding LC3B, we used bacterial surface display in combination with fluorescence-activated cell sorting (FACS). Following Hwang *et al*., we expressed ∼500,000 36-mer peptides spanning the human proteome on the cell surface of *Escherichia coli* via fusion to <u>e</u>nhanced <u>c</u>ircularly <u>p</u>ermuted Omp<u>X</u> (eCPX) (Rice *et al*. 2006; Larman *et al*. 2011; Hwang *et al*. 2022). To compensate for the low micromolar affinity of typical SLiM interactions, and to better mimic the avid interactions enabled by the high local concentration of LC3B conjugated to autophagosomal membranes in cells (Zaffagnini and Martens 2016), we bound N-terminally biotinylated LC3B to tetravalent streptavidin-phycoerythrin (SA-PE). Using FACS, we quantified peptide expression levels using an allophycocyanin (APC)-conjugated antibody against an encoded N-terminal FLAG tag. Cells displaying peptides that bound to LC3B were identified and isolated based on their high SA-PE-to-APC ratio (**Figure 1A**). We validated our screening approach using well-characterized LC3B-binding peptides of various affinities (**Supplementary Figure 1**). Following five rounds of positive selection and one round of counterselection against non-specific binding to SA-PE (**Figure 1B**), cells in the binding gate were enriched ∼80-fold, from ∼0.7% of the naïve library to ∼57% of all peptide-expressing cells (**Supplementary Figure 2**).

**Figure 1.**
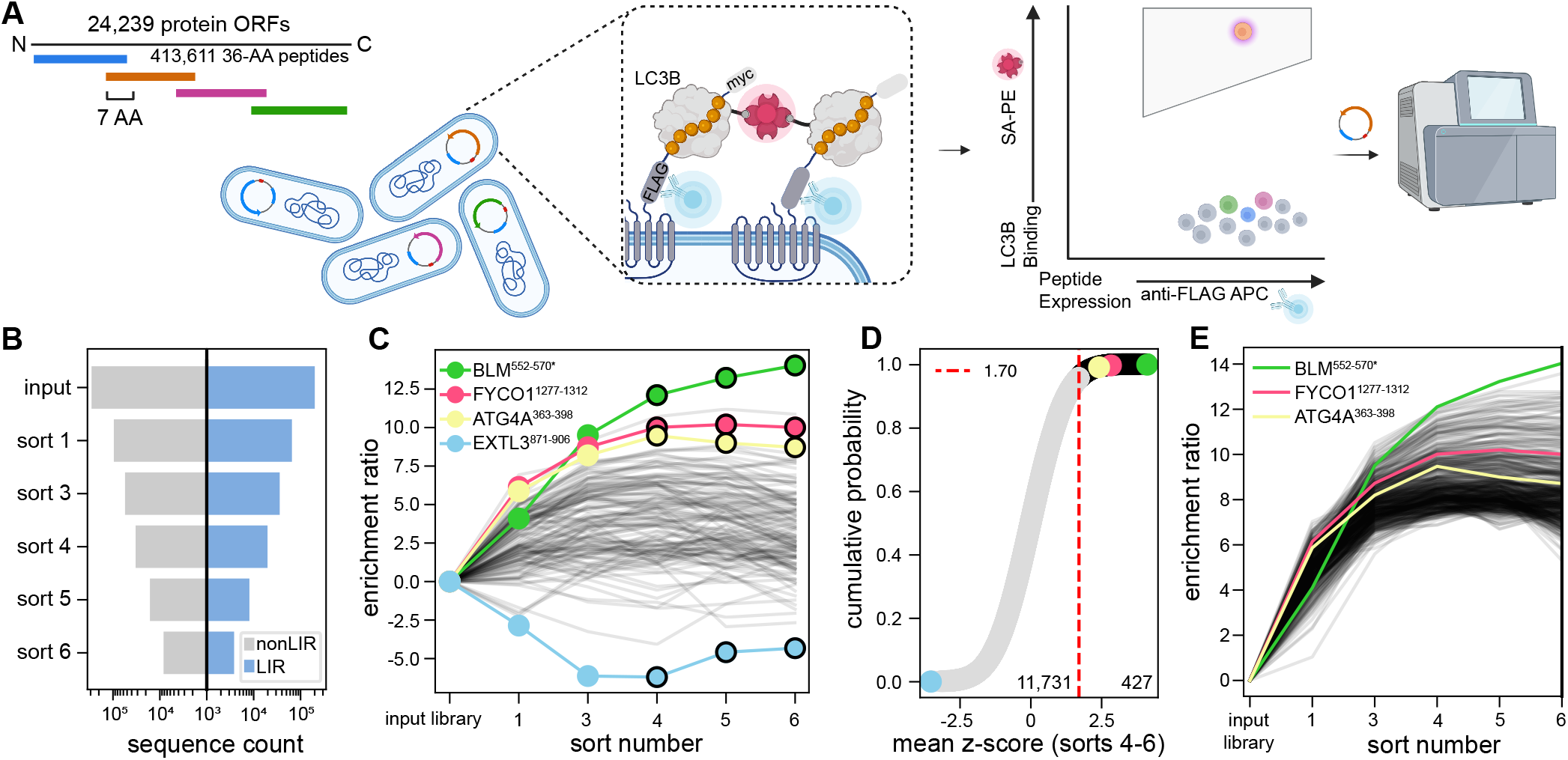
Human peptidome library screening enriches known and novel LC3B-binding peptides. **(A)** Schematic of FACS-based bacterial display enrichment screening. Peptides, 36 amino acids in length, that tile the human proteome were expressed as FLAG-tagged fusions to eCPX on the surface of *E. coli*. Expression was detected using the fluorescence signal from an allophycocyanin (APC)-conjugated anti-FLAG antibody. Binding to LC3B (grey), which was tetramerized through binding to streptavidin conjugated to phycoerythrin (SA-PE, pink), was detected using PE fluorescence. Peptide-expressing cells were sorted based on APC and PE fluorescence and sequenced. Created with biorender (https://biorender.com/7v3ijfa). **(B)** Diverging bar chart plotting the number of unique sequences detected in each round of sorting that lacked (grey, left axis) or contained (right, blue) a LIR motif. Sort 2, a negative sort used to eliminate peptides nonspecifically bound to SA-PE, was not sequenced. **(C)** Peptide enrichment profiles across enrichment sorts. Black lines show trajectories for a random subset (0.1%) of all peptides that persisted to sort 6. Overlaid are the enrichment profiles for the best-enriching peptide (BLM^552-570*^), positive controls (FYCO1^1277-1312^ and ATG4A^363-398^), and the worst-enriching peptide that reached sort 6 (EXTL3^871-906^), colored green, pink, yellow, and blue, respectively. Peptide sequences are detailed in **Supplementary Table 1**. **(D)** Cumulative density function plot of the mean z-scores across sorts 4, 5, and 6. Mean z-scores for the 427 peptides of 12,158 that surpassed the threshold of 1.70 that were used to define the HC-set are colored in black, and select peptides are colored as in panel C. **(E)** Peptide enrichment profiles for the 427 peptides in the HC-set (black), with select peptides colored as in panels C-D.

In all, 12,158 peptides derived from 5,578 unique proteins were identified in the final round of library sorting, with individual peptides displaying a wide range of behaviors as assessed by their calculated enrichment ratio (ER) (Rubin *et al*. 2017) across successive sorts (**Figure 1C**, see Methods). To define a high-confidence set (HC-set) of LC3B binders for further analysis, we selected peptides with a mean ER at least 1.70 standard deviations (z-score ≥ 1.70, n = 427 peptides) above the mean ER calculated over the final three rounds of sorting (**Figure 1D-E**). This criterion captured four of the seven peptides in our input library that were reported to bind LC3B with dissociation constants below 5 µM, as compiled in the LIRcentral database (Chatzichristofi *et al*. 2023) (**Supplementary Figure 3A**). In contrast, peptides in our input library annotated by LIRcentral as non-binding (K_D_ > 50 µM) were strictly excluded from this group and were progressively depleted during sorting (**Supplementary Figure 3B**). Beyond these known interactions, this HC-set included peptides from sixteen proteins previously reported to co-immunoprecipitate with LC3B (Stark *et al*. 2006), most of which lacked mapped interaction sites (**Figure 2A**). We tested four such peptides (MAP1B^823-858^, ATG4A^363-398^, SCYL1^640-675^, and HEAT3^377-412^) for binding to monomeric LC3B by bio-layer interferometry (BLI) and observed robust binding with dissociation constants ranging from approximately 0.5 to 10 µM (representative plots in **Figure 2B**, all data in **Supplementary Table 1**).

**Figure 2.**
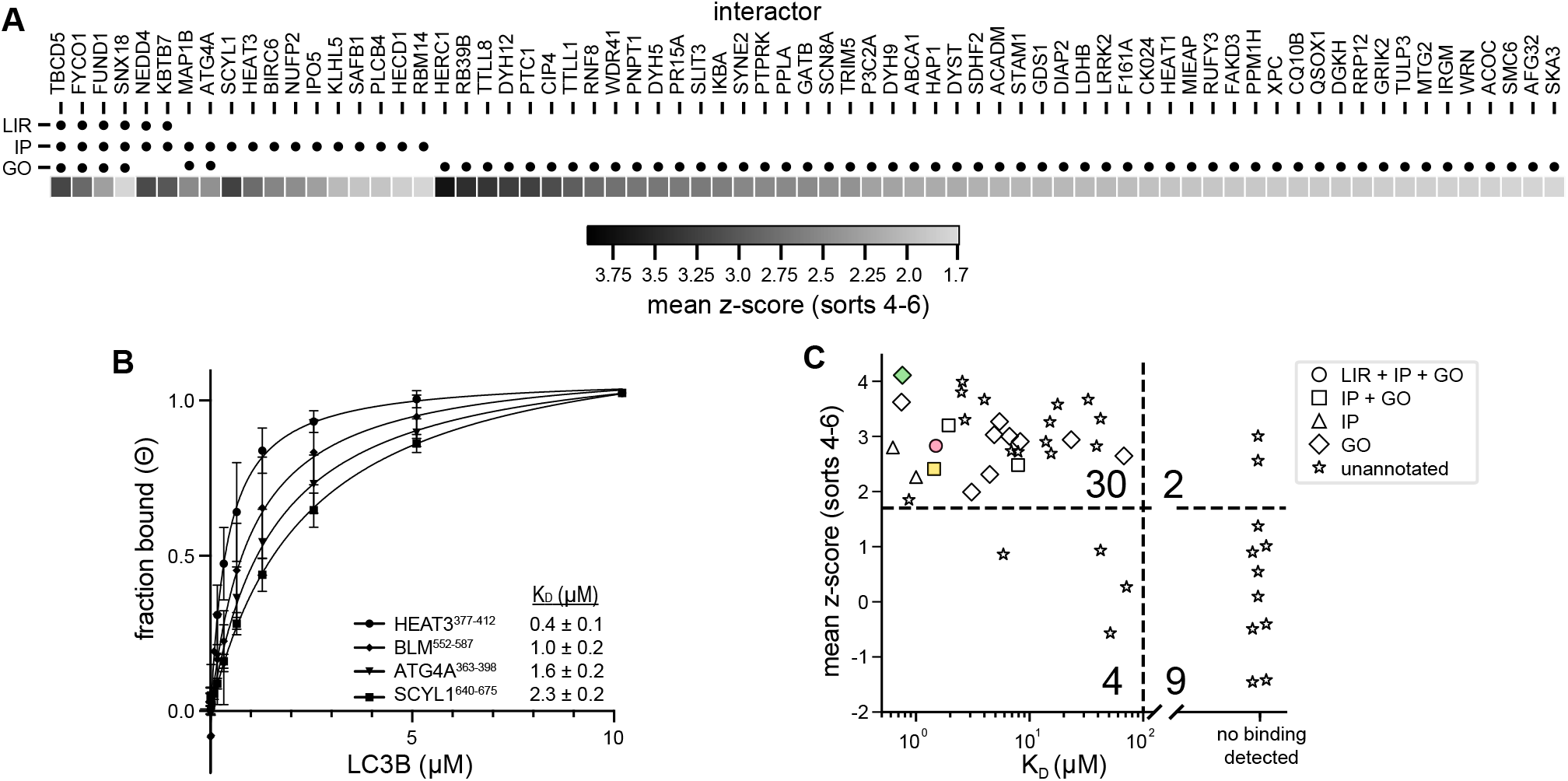
Highly enriched peptides share annotations with LC3B and bind LC3B tightly. **(A)** Peptides in the HC-set, annotated with various orthogonal lines of evidence consistent with binding to LC3B. For each protein, circles indicate: LIR - experimentally validated LIR motifs annotated in LIRcentral (Chatzichristofi *et al*., 2023); IP - proteins co-purified with LC3B, from BioGRID (Stark *et al*., 2006); GO – peptides with GO-annotations shared with LC3B, from UniProt (The UniProt Consortium, 2025), with heatmap of enrichment z-score plotted at bottom. **(B)** Binding to monomeric LC3B assayed by BLI for peptides ATG4A^363-398^ (VPPAKPEVTTTGAEFIDSTEQLEEFDLEEDFEILSV), SCYL1^640-675^ (TADRWDDEDWGSLEQEAESVLAQQDDWSTGGQVSRA), HEAT3^377-412^ (EDPSDDEWEELSSSDESDAFMENSFSECGGQLFSPL), and BLM^552-587^ (DIDNFDIDDFDDDDDWEDIMHNLAASKSSTAAYQPI). Assay performed in triplicate, with error bars indicating standard deviation. Data fit to a standard binding isotherm: 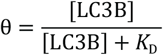 **(C)** Relationship between mean z-score over rounds 4-6 and affinity for monomeric LC3B as determined by BLI. Three peptides are colored as in **Figure 1C**. Shapes indicate five classes of peptides, as annotated in panel A. Dotted lines mark thresholds delineating high vs. low z-scores and binding vs. non-binding, with the number of peptides in each quadrant listed. Enrichment of binders among high z-score peptides was found to be statistically significant, as assessed using Fisher’s exact test (p < 0.001).

### Strongly enriched peptides include novel candidate LC3B interactors

To identify peptide hits from our screen most likely to represent biologically relevant LC3B interaction partners, we selected proteins with shared Gene Ontology (GO) annotations (The Gene Ontology Consortium *et al*. 2023) with LC3B that contained peptides in our HC-set (**Figure 2A, Supplementary Figure 4**). From this list, we tested LC3B for binding to two candidate peptides derived from HERC1^3075-3110^ and LRRK2^858-893^, noting that these proteins are reported to impact autophagy (Montes-Fernández *et al*. 2020; Pérez-Villegas *et al*. 2020; Park *et al*. 2016; Roosen and Cookson 2016; Madureira *et al*. 2020; Boecker and Holzbaur 2021), but have not been shown to directly interact with LC3B. Each peptide bound with a dissociation constant in the low micromolar range (**Supplementary Table 1**).

We additionally identified peptide hits without reported links to autophagy or LC3B. Many of these peptides bound LC3B with moderate affinity (K_D_ ≤ 50 µM) and thus represent candidates for new LC3B-interacting proteins (**Supplementary Table 1**). In total, we determined monomeric LC3B binding affinities for 45 peptides spanning varying degrees of enrichment in our sort and found that the z-score cutoff of 1.70 used to define our HC-set effectively distinguished binders (K_D_ < 50 µM) from non-binders (K_D_ > 60 µM) (**Figure 2C, Supplementary Table 2**). Two peptides with z-scores greater than 1.70 in the screen (from PAR12 and KBTB7) did not bind detectably in the BLI assay; this may be due to the higher avidity of the interaction with LC3B in the bacterial display format.

### Enriched LIR-containing sequences exhibit a preference for Trp in the X_0_ position, preceded by acidic residues

To identify sequence features associated with binding, we generated a sequence logo (Schneider and Stephens 1990; O’Shea *et al*. 2013) for peptides in the HC-set that contained a LIR motif (see Methods). This logo highlighted enrichment of tryptophan in the first position (X_0_), over tyrosine and phenylalanine, and a preference for acidic residues in positions X_-1_, X_-2_, and X_-3_, as well as for glutamate in the X_1_ position of the core LIR motif and at the C-terminal X_7_ position (**Figure 3A**). Consistent with this logo capturing features related to binding affinity, a synthetic peptide defined by the consensus sequence observed in the logo, pCONS_LIR_, bound to LC3B with a dissociation constant of ∼60 nM (**Figure 3B**). This affinity is ∼30-fold tighter than a well-studied peptide from FYCO1^1277-1312^ (Olsvik *et al*. 2015; Cheng *et al*. 2016), ∼25-fold tighter than a chimeric peptide designed to target α-synuclein for autophagic degradation (Tong *et al*. 2023), and is comparable in affinity to the tightest known LC3B-binding peptide, ANK2^1578-1613^ (Li *et al*. 2018).

**Figure 3.**
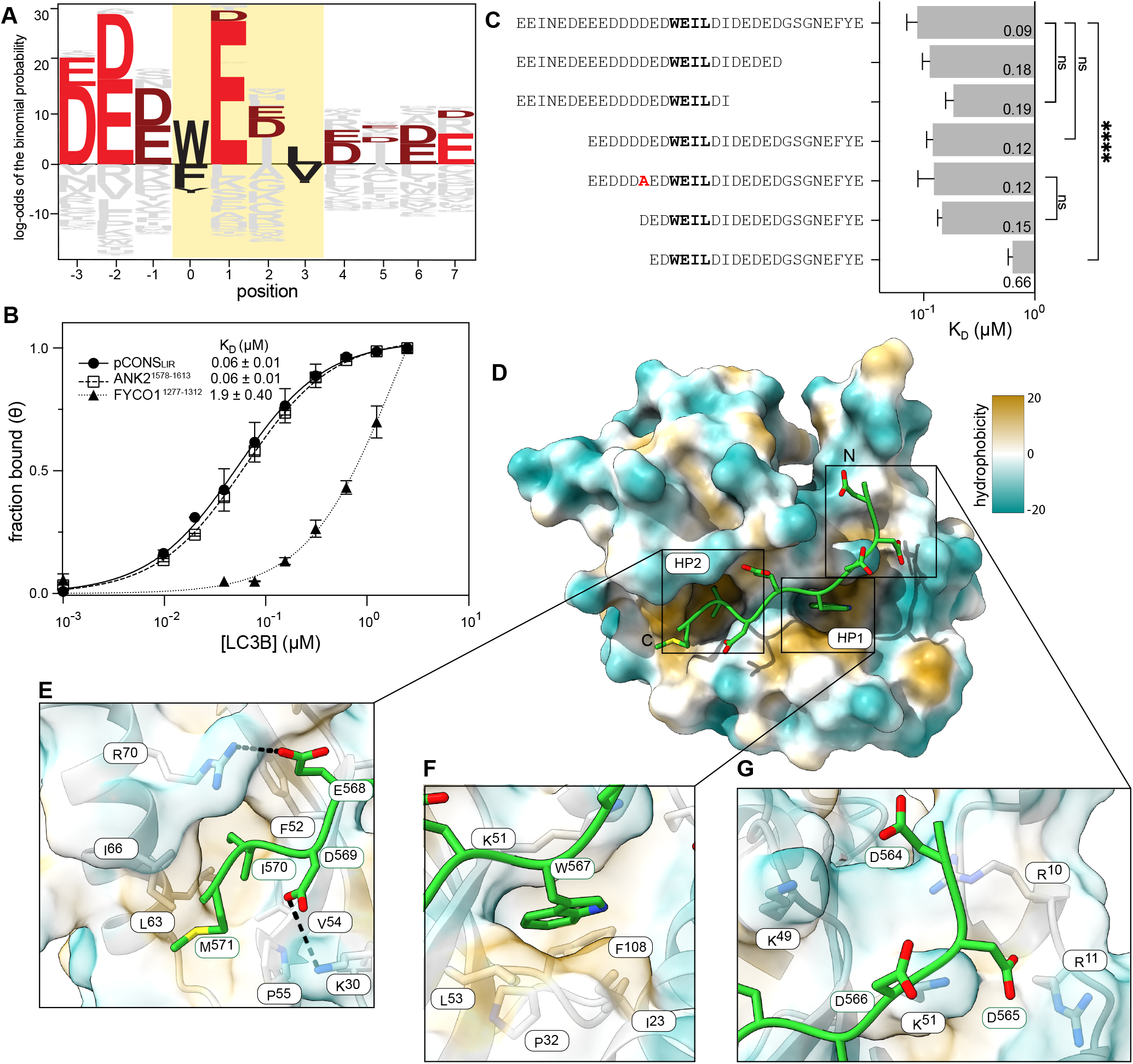
Highly enriched peptides feature W-type core LIR motif flanked by acidic residues. **(A)** Sequence logo derived from analysis of the HC-set peptides (see Methods), and plotted using Logomaker (Tareen and Kinney, 2020). Acidic residues are colored maroon and those colored red are significantly enriched compared to the input library. Other residues are colored gray with [FWY]_0_ and [LVI]_3_ in black. The core LIR is highlighted in yellow. **(B)** BLI measurements of binding affinity between LC3B and peptides pCONS_LIR_ (EEEVEEKEEEDDDEEWEILDIEEGSDSEQKLISE), ANK2^1578-1613^ (VQSSRSERGLVEEEWVIVSDEEIEEARQKAPLEITE), and FYCO1^1277-1312^ (DAVFDIITDEELCQIQESGSSLPETPTETDSLDPNA). Error bars report the standard deviation of two or more technical replicates. Data fit to a standard binding isotherm, as in **Figure 2B**. **(C)** Bar chart plotting the mean binding affinities (K_D_) of sequential truncations of pCONSw measured via BLI, with error bars reporting standard error of the mean (n = 3). Brackets report statistical significance of pairwise t-tests (\*\*\*\**p* ≤ 0.0001; ns not significant). Values indicated in the bars are K_D_ values in μM. **(D)** Structure of BLM^554-571^ bound to LC3B, resolved to 2.0 Å resolution (see **Supplementary Table 4**). Here, and in panels E-G, the BLM peptide is displayed in stick representation (green backbone) and the LC3B surface is colored by hydrophobicity, as computed by ChimeraX (Meng *et al*., 2023). Hydrophobic pockets HP1 and HP2, and the N-terminal flanking residues, are indicated. **(E)** Inset highlighting contact between BLM E568 and LC3B R70, and BLM D569 and LC3B K30, near HP2 (black dashed lines). LC3B residues that form HP2 are annotated and shown in stick representation. **(F)** Inset highlighting docking of BLM W567 in LC3B HP1, with residues forming that pocket annotated and shown in stick representation. **(G)** Inset highlighting interactions between BLM N-terminal acidic residues D564 and D566 with R11, K49, and K51 of LC3B. Putative contacts within 4.5 Å marked with dashed black lines.

To test the role of the hydrophobic residue in position X_0_, we partitioned the sequence alignments based on the W/F/Y residue identity and generated three new peptides (pCONS_W_, pCONS_F_, and pCONS_Y_) based on the consensus sequences (**Supplementary Figure 5**). Because Y-type LIR sequences were frequently located near the C-terminus in our HC-set, we lacked sufficient sequence diversity to generate a logo with C-terminal flanking sequences for this region, and the LIR motif of pCONS_Y_ was placed at the C-terminus of the peptide to reflect this positional bias. Peptide pCONS_W_ bound to LC3B with similar affinity (K_D_∼90 nM) as peptide pCONS_LIR_ (**Supplementary Table 3**). The affinities of pCONS_F_ and pCONS_Y_ were both weaker (K_D_∼400 nM, K_D_∼10 µM, respectively) (**Supplementary Table 3**), consistent with tryptophan in the first position (X_0_) favoring binding.

We next tested the contributions of the highly enriched N- and C-terminal acidic residues in the context of the tightly binding pCONS_W_ peptide. Truncation of C-terminal residues beyond position X_5_ had modest effects on binding, but removing residues N-terminal to position X_-2_ weakened affinity ∼7-fold (**Figure 3C, Supplementary Table 3**). Most of this reduced affinity could be restored by reintroducing either a single N-terminal Asp or a 6-residue EDDDDA sequence that lacked this Asp (**Figure 3C, Supplementary Table 3**). These data support an important role for flanking N-terminal acidic residues in facilitating high-affinity binding to LC3B, without a strict requirement for a specific residue at a specific site. Consistent with the key determinants of binding being proximal to the LIR, removing 7 residues from either the N- or C-terminus of this peptide did not significantly impact affinity (**Figure 3C, Supplementary Table 3**).

Given the efficacy of this approach in predicting tightly binding peptides, we next evaluated whether a related method could distinguish LC3B-binding from non-binding peptides that contained LIR motifs. Briefly, we assembled a test set of 79 binders and 56 non-binders as annotated in LIRcentral (Chatzichristofi *et al*. 2023) and compared the discriminatory performance of the previously reported iLIR model (Kalvari *et al*. 2014) with new PSSM-based models derived from our screening data (see Methods).

We also constructed higher-order random forest classifiers using our screening data that aimed to capture potential residue–residue coupling effects. On this test set (see Methods), no model performed better than the iLIR model (**Supplementary Figure 6**). Evaluating the models using held-out screening data gave auROC = 0.87 for recognizing LIR peptides with z-score > 2.3, but poor performance for less enriched peptides (**Supplementary Figure 7**), suggesting that the test set and/or screening data may be too limited, sparse, or noisy, or that key binding determinants depend on higher-order structure that our models fail to capture.

### Acidic residues N-terminal of the core LIR directly contribute to LC3B engagement and binding affinity

Many of our top-scoring peptides exhibited features common to both pCONS_LIR_ and pCONS_W_, namely the presence of Trp in X_0_, Glu in X_1_, and multiple N-terminal acidic residues. Indeed, the top-scoring peptide contained these features, and bound tightly to LC3B (K_D_∼0.7 µM; **Supplementary Table 1**). Many previously reported high-affinity LC3B binders, including FYCO1^1276-1288^, ANK2^1588-1613^, and ANK3^1985-2010^, use an acidic residue in position X_7_ of the C-terminal extension to make an affinity-enhancing contact to LC3B residue R70 (Olsvik *et al*., 2015; Cheng *et al*., 2016; Li *et al*., 2018) (**Supplementary Figure 8**), yet our top scoring peptide, derived from the DNA helicase BLM, lacks such a residue. This observation prompted us to investigate how it achieved comparable binding.

We determined the structure of LC3B fused to BLM^554-571^ at 2.0 Å resolution using X-ray crystallography (**Figure 3D-G, Supplementary Table 4**, pdb 9p3e). BLM^554-571^ engaged the LDS of LC3B as expected, and the interactions between the peptide and LC3B were consistent with those observed for other canonical LIR motifs: the aromatic W567 (X_0_) and hydrophobic residue I570 (X_3_) docked in HP1 and HP2, respectively (**Figure 3E-F**). Additionally, we observed an intermolecular β-sheet between the LC3B residues 51-53 and BLM residues 568-570, which occupy the 1-3 positions of the [FWY]_0_-X_1_-X_2_-[LVI]_3_ LIR motif. Notably, the sidechain of BLM E568 (X_1_) contacted LC3B R70 (**Figure 3E**), seemingly substituting for the contacts often found between R70 and glutamates C-terminal to the core LIR (**Supplementary Figure 8**). Acidic residues N-terminal to the LIR, D564-D566 (X_-3_ – X_-1_), were in proximity to positively charged residues R10, R11, K49, and K51 of LC3B (**Figure 3G**), though they adopted multiple conformations as analyzed by ensemble refinement (Burnley *et al*. 2012) (**Supplementary Figure 9**, see Methods). Together, these observations indicate that acidic residues N-terminal to the core LIR contribute directly to LC3B engagement and binding affinity.

### Highly enriched peptides lacking a canonical LIR can bind LC3B

We next asked whether sequences from our HC-set that lacked a canonical LIR could also bind LC3B, focusing on candidates with residues we predicted could engage the HP1 and HP2 pockets. We specifically examined peptides containing a motif we termed LIR+ ([FWY]_0_-X_1_-X_2_-[FWY]_3_), as Li *et al*. had previously reported an X-ray crystal structure showing a peptide bound to LC3B with aromatic residues W and F engaged with the hydrophobic pockets (Li *et al*., 2018). To this end, we assessed LC3B binding for three peptides in our HC-set bearing LIR+ sequences: (1) DYH12^422-445^*, which contains two LIR+ sequences; (2) PAR11^8-37^, which contains a single LIR+ sequence; and (3) CTSL2^233-262*^, which contains both a canonical LIR and a LIR+ motif. We found that each peptide bound LC3B with sub-20 µM affinity as measured by bio-layer interferometry (BLI). We then introduced alanine substitutions into each LIR+ sequence and observed reduced affinity for LC3B in each peptide, consistent with a role for the LIR+ motif in supporting LC3B association (**Figure 4A**). Notably, in CTSL2^233-262*^, mutation of the LIR+ sequence ^252^WEVF^255^ to AAAA strongly reduced binding, whereas analogous alanine substitutions in the canonical ^258^YYFI^261^ LIR motif had a modest effect, consistent with the LIR+ site serving as the dominant binding determinant. Supporting this interpretation, an AlphaFold3 (Abramson *et al*., 2024) prediction of a CTSL2^233-262*^•LC3B complex structure showed the LIR+ motif, and not the LIR motif, engaging the LC3B LDS (**Figure 4B**). To compare the WEVF motif binding mode to canonical LIR binding at the LDS, we predicted the structure of CTSL2^233-262*^•LC3B with the WEVF motif mutated to WEVI. The two structures were very similar, with a small backbone shift of 2 Å at the mutated site that accommodates the larger aromatic residue in the LIR+ motif.

**Figure 4.**
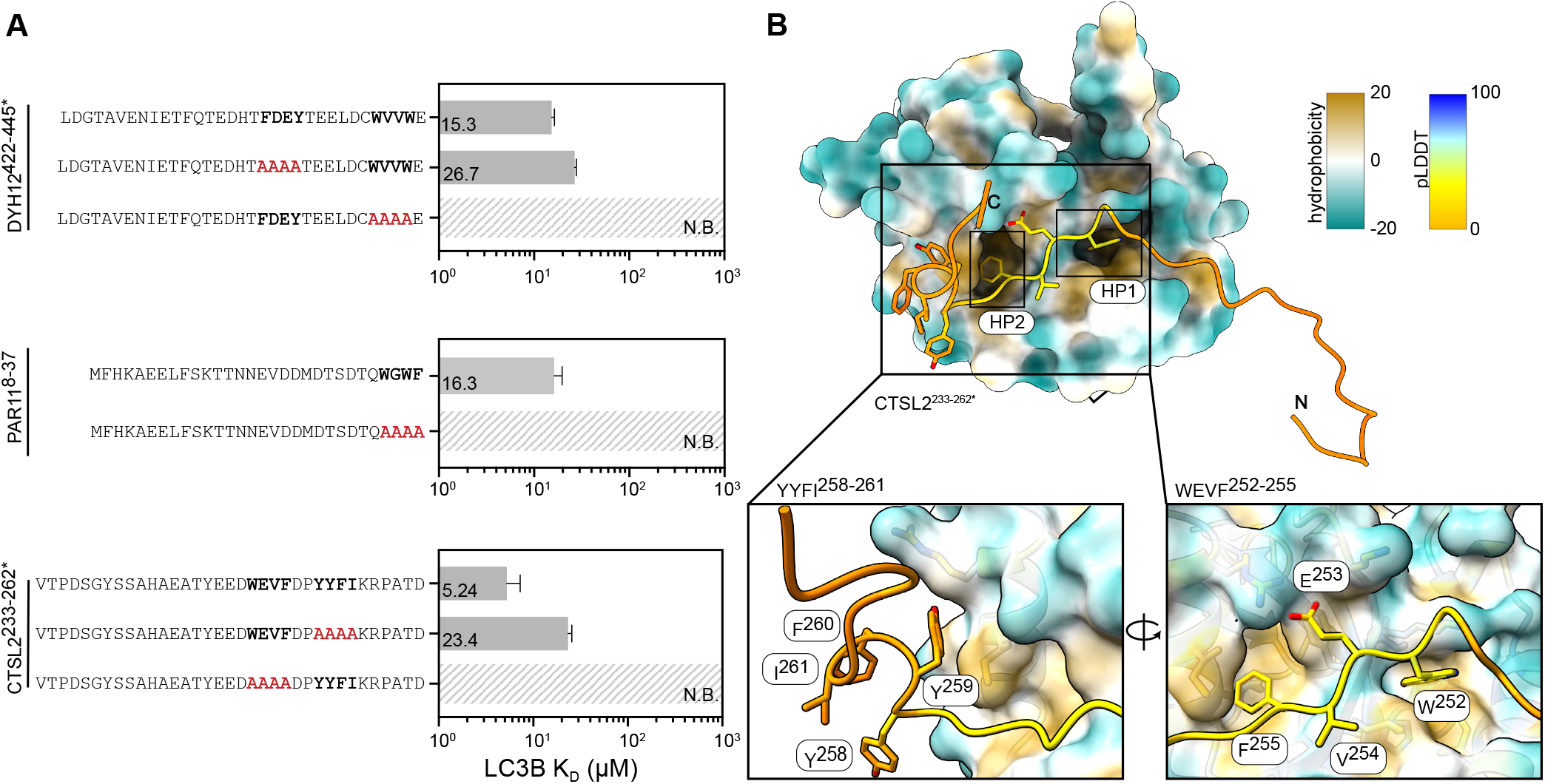
LIR+ motifs support binding to LC3B. **(A)** Affinities of LC3B for peptides from DYH12^422-445^*, PAR11^8-37^, and CTSL2^233-262*^, as measured by BLI, plotted as a bar chart (mean ± s.e.m, *n*≥3). Candidate binding sites that were mutated to Ala are red in the mutated sequence. Values indicated in the bars are K_D_ values in μM. No detectable binding, up to 40 µM LC3B, is indicated with N.B. **(B)** AlphaFold3 (Abramson *et al*., 2024) structural prediction of CTSL2^233-262*^ (colored by pLDDT) in complex with LC3B (colored by hydrophobicity). Insets: canonical LIR YYFI^258-261^ is not predicted to engage the LDS (left), in contrast to WEVF^252-255^ (right).

Considering the apparent binding efficacy of such LIR+ sequences, we assessed the relative frequency of canonical LIR, LIR+, and related motifs in our input library and in our HC-set. We found strong evidence that canonical LIR motifs ([FWY]xx[LVI]) are enriched in the HC-set relative to their frequency in the input library, whereas LIR-related motifs ([WFY]xx[WFY], [LVI]xx[FWY], [LVI]xx[LVI]) are not (**Supplementary Figure 10**), indicating that although LIR-related motifs can support LC3B binding in certain sequence contexts, they are not sufficiently strong determinants to drive enrichment in our screen.

### Common mutations to the LIR docking site of LC3B modulate but do not ablate peptide binding

LC3B with mutations F52A and L53A in the LDS, here called LC3B LDS*, is commonly used to assess binding at the LDS, with these substitutions in HP2 and HP1 presumed to disrupt peptide binding at this interface (**Figure 5A**) (Behrends *et al*. 2010; Kraft *et al*. 2014; Qiu *et al*. 2017; Skytte Rasmussen *et al*. 2017; Marshall *et al*. 2019). Using LC3B LDS*, we performed three additional bacterial-display sorts with the round-5-enriched library as input (**Supplementary Figure 11**, see Methods). The well-studied peptide FYCO1^1277-1312^ was depleted over the three rounds of LDS* sorting, consistent with its canonical LDS-dependent binding mechanism. Similar behavior was observed for other known LC3B-binding peptides, including FUND^11-36^, NEDD4^681-716^, and ATG4A^363-398^ (**Figure 5B**). Unexpectedly, several other peptides were further enriched over these sorts (**Figure 5B**), consistent with increased affinity for the altered LC3B LDS* interface.

**Figure 5.**
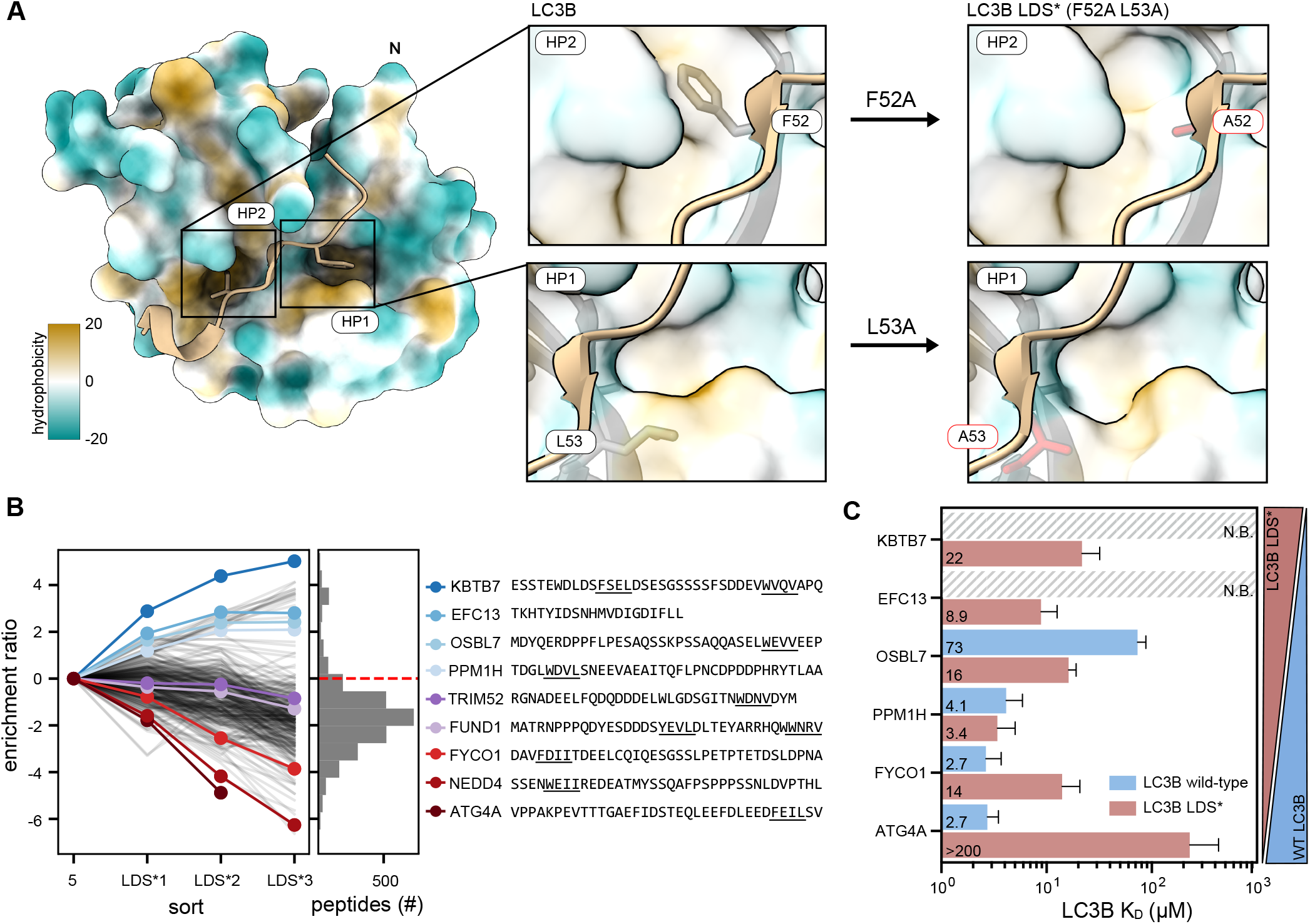
Mutations at the LC3B LIR docking site alter LIR-peptide binding specificity. **(A)** Structure of LC3B bound to FYCO1^1276-1288^ LIR peptide (PDB 5d94) (Olsvik *et al*. 2015) shown as a surface representation colored by hydrophobicity. Insets depict hydrophobic pockets formed by residues F52, L53 (stick representation) in wild-type LC3B (left), or a predicted surface of the LDS* mutant (right), with each alanine substitution colored red. **(B)** Enrichment profiles across three rounds of the LC3B LDS* enrichment sort, with nine peptides spanning the range of observed profiles colored and sequences provided. Four peptides that were enriched and bound with measurable affinity to LC3B LDS* are in blue. Known, canonical LC3B LDS binders that were depleted in the LDS* sort are colored purple and red. Black lines indicate a random sample of 10% of peptides that persisted through LC3B LDS* sort 3. The histogram shows the frequencies of peptides with a given enrichment ratio in LDS* sort 3, with the dashed red line marking an enrichment ratio of 0. **(C)** Affinities of LC3B (blue) and LC3B LDS* (light red) for peptides FYCO1^1277-1312^ and ATG4A^363-398^ and for four peptides found to enrich across the LDS* sorts: PPM1H^436-471^, OSBL7^1-36^, EFC13^262-278*^, KBTB7^639-674^. Values indicated in the bars are K_D_ values in μM. Error bars indicate standard error of the mean across three or more replicates. Hashed bars mark peptides without detectable binding at the highest measured LC3B concentration (40 µM).

To investigate this further, we selected six peptides spanning the enrichment patterns observed in the LDS* sorts and quantified their binding to wild-type LC3B or LC3B LDS* proteins using BLI. The results revealed diverse binding behaviors: some peptides bound more tightly to wild-type LC3B (*e*.*g*., ATG4A^363-398^), whereas others exhibited strong selectivity towards the LDS* mutant (*e*.*g*., KBTB7^639-674^) (**Figure 5C**). Notably, structural modeling indicated that although the LDS* mutations alter the binding interface, the hydrophobic pockets persist (**Figure 5A**). Therefore, we propose that, rather than abolishing LDS-mediated interactions entirely, the F52A/L53A substitutions subtly remodel the docking surface, creating an alternative hydrophobic interface that modulates peptide binding in a sequence-dependent manner.

## DISCUSSION

LC3B and other hAtg8 paralogs often interact with binding partners via the LIR motif [FWY]_0_-X_1_-X_2_-[LVI]_3_, an information-poor SLiM that can mediate critical interactions in all major stages of bulk and selective autophagy (Rogov *et al*., 2023), as well as in non-autophagic processes such as LC3-associated phagocytosis (Florey and Overholtzer 2012) and LC3-associated endocytosis (Heckmann *et al*., 2019). Here, we used high-throughput bacterial-surface-display screening to broadly assess the LC3B-binding potential of peptides derived from the human proteome, and to define sequence determinants of LC3B recognition. We focused our analysis on a set of strongly enriched peptides (*i*.*e*., the HC-set), recognizing that we would miss many authentic but weakly interacting partners. Within the HC-set, we observed roughly threefold enrichment of peptides bearing traditional LIR sequences. Many peptides contained multiple LIR motifs, which may enhance apparent affinity through avid interactions with LC3B oligomerized through the tetravalent streptavidin linkage used in our assay, or through membrane conjugation in cells.

Overall, the vast majority (∼90%) of peptides in the HC-set contained a canonical LIR or a related sequence bearing hydrophobic or aromatic residues positioned to plausibly interact with hydrophobic pockets HP1 and HP2 on LC3B. We biochemically characterized one such motif [WFY]_0_-X_1_-X_2_-[WFY]_3_, which we termed “LIR+”, finding that it supported robust LC3B binding. Although a small number of LC3B-binding peptides bearing this LIR+ motif have been reported (∼1% of cataloged hAtg8-binding peptides) (Li *et al*., 2018; Chatzichristofi *et al*., 2023), their prevalence appears underappreciated given their relatively high frequency in the HC-set (**Supplementary Figure 10**), leading to the refined motif [FWY]xx[LVIFWY]. Further, the binding of LIR+ peptides observed here suggests that other motif variants, such as [ILV]_0_-X_1_-X_2_-[WFY]_3_ or [ILV]_0_-X_1_-X_2_-[ILV]_3_, may also engage LC3B and, consistent with this notion, such sequences were prevalent in our HC-set (**Supplementary Figure 10**).

The remaining peptides lacking any LIR-like motifs (∼10%) partitioned between those in which the two hydrophobic residues were separated by either 3 or 4 acidic amino acids (3 peptides), similar to the LC3B-binding “<u>W</u>DDE<u>W</u>” motif in SNX18 (Knævelsrud *et al*., 2013), those bearing the sequence “HPQ” (3 peptides), which is reported to bind to streptavidin (Weber *et al*., 1992; Katz 1995), and a set of peptides lacking linear motifs identifiable through expert-guided inspection or the automated motif discovery tool XSTREME (Grant and Bailey, 2021).

A sequence logo-based analysis of LIR-bearing peptides in our HC-set revealed enrichment of acidic residues, and biochemical measurements confirmed that acidic residues N-terminal to the core LIR enhance binding affinity without strict positional dependence. These observations complement and expand on previous studies of individual peptides derived from FYCO1 and p62/SQSTM1, for which LC3B binding is supported by N-terminal acidic residues (Pankiv *et al*., 2007). They are also aligned with a reported affinity-enhancing role for phosphorylated serine and threonine residues in the N-terminal flanking sequence of the core LIR motif (Richter *et al*., 2016; Wirth *et al*., 2021; Kliche *et al*., 2023). Indeed, given the impact of N-terminal acidic residues, it is plausible that phosphorylation of target proteins could dynamically modulate LIR•LC3B binding affinity, effectively linking kinase-dependent environmental signaling cascades to the regulation of autophagic cargo selection (Birgisdottir *et al*., 2013; Kliche and Ivarsson 2022; Rogov *et al*., 2023).

LIR databases catalog experimentally validated motifs (Chatzichristofi *et al*., 2023) and have been used to define a six-residue PSSM model for predicting binders to hAtg8 proteins and other Atg8 orthologs (Kalvari *et al*., 2014). Guided by sequence information from our screening data, we designed a 34-residue synthetic peptide with an affinity comparable to the strongest known interaction partners (**Figure 3B, Supplementary Table 3**). The high-affinity pCONS_LIR_ peptide is characterized by an extended negatively charged N-terminal flanking sequence. Notably, the peptides exhibiting the strongest enrichment in our screen (*i*.*e*., z-score ≥ 2.3) and those described in the iLIR set of known binders shared this prominent enrichment of acidic N-terminal residues, as evidenced by these sequences co-clustering in UMAP space (**Supplementary Figure 6D**). However, peptides lacking these specific features can also bind LC3B, and others with similar sequence features do not bind LC3B, underscoring the complex relationship between peptide sequence and LC3B recognition that was not fully captured by the PSSM-based classifiers or by the random forest classifiers we deployed (**Supplementary Figures 6-7**).

Recently, the Vierstra group reported an alternate binding interface on Atg8 homologs (hAtg8) that they termed the <u>U</u>biquitin-interaction-motif <u>D</u>ocking <u>S</u>ite (UDS). This interface, which occurs on the face opposite that of the LIR docking site, is reported to interact with an alternative SLiM known as the <u>U</u>biquitin interacting <u>m</u>otif (UIM), with the sequence Ψ-ζ-X-A-Ψ-X-X-S, where Ψ, ζ, and X denote small hydrophobic, hydrophilic, and any amino acids, respectively (Marshall *et al*., 2019). Peptides bearing this motif were present in our input library (2,452 peptides) yet only 4 were found in our HC-set, indicating that if an LC3B UDS exists, it was either inaccessible in our assay format or did not support sufficiently high-affinity binding.

Notably, we did not observe enrichment for UIM-bearing peptides even when screening in the context of LC3B constructs bearing mutations at the LIR docking site (LDS*). Instead, screening against LC3B LDS* revealed that peptides bearing canonical LIR sequences exhibited altered affinity for the mutant LC3B domain (**Figure 5**), consistent with these often-utilized mutations changing specificity but not ablating binding, contrary to what has been assumed (Behrends *et al*., 2010; Kraft *et al*., 2014; Qiu *et al*., 2017; Skytte *et al*., 2017; Marshall *et al*., 2019).

Together, the LC3B and LC3B LDS* bacterial display datasets presented here constitute a rich resource for the study of SLiMs and autophagy. These datasets reveal preferred residues flanking the canonical LIR motif, expand the repertoire of LC3B-binding sequences, and highlight important caveats in using the LC3B F52A L53A LDS* mutant to disrupt LDS-dependent interactions. Although peptides in our bacterial display system cannot fully capture the complexity of LC3B binding in cells— where SLiM accessibility, subcellular localization, and post-translational modifications may play key roles—our results provide a quantitative and structural framework for understanding LC3B recognition. Moreover, this work nominates new candidate LC3B partners for validation in cellular and physiological contexts and establishes a generalizable framework for high-throughput analysis of SLiM-peptide interactions with Atg8-family proteins.

## MATERIALS AND METHODS

### Expression and bacterial display vector construction

#### LC3B expression vectors

Coding sequences for LC3B and the LDS* mutant (F52A, L53A) were cloned into plasmid pDW363 (Tsao *et al*., 1996) downstream of an N-terminal Biotin Acceptor Peptide (BAP) and 6×His tag (GLNDIFEAQKIEWHE-DTGGSS-HHHHHH-GSGSG-[coding sequence]). The resulting plasmids (pl_ JD239, pl_JD362) also expressed the BirA biotin ligase for *in vivo* biotinylation of the BAP-fused LC3B. For BLI experiments, the BAP tag was removed to yield 6×His-tagged, non-biotinylated variants pl_JD361 (LC3B wild-type) and pl_JD363 (LC3B LDS*) (M-GSS-HHHHHH-GSGSG-[coding sequence]). For crystallization of LC3B bound to the BLM^554-571^ peptide, DNA constructs were designed to fuse the sequence DNFDIDDFDDDDD<u>WEDI</u>M N-terminal to LC3B, separated by a two-residue GS linker, and cloned into a pGEX vector containing an N-terminal GST and 3C cleavage site using Gibson assembly (Gibson *et al*., 2009), resulting in plasmid pl_JD371.

#### Peptide expression vectors

The coding sequence of each peptide was synthesized either as a gene block (IDT) or was encoded on primers (Eton Bio), each used to insert this coding sequence into a modified pDW363 vector using Gibson (Gibson *et al*., 2009) or ‘round-the-horn’ insertional cloning (Liu and Naismith, 2008). The resulting products fused each peptide to the C-terminus of a BAP–6×His–SUMO tag via a short (SG)_2_ linker.

#### Bacterial display vectors

The human peptidome bacterial display library is described by Hwang *et al*., and the peptide-encoding DNA was originally provided by the Elledge laboratory based on work of Larman *et al*. Briefly, each peptide is fused to the C-terminus of enhanced circularly permuted *E. coli* OmpX (eCPX) (Rice *et al*., 2006), with the peptide flanked by an N-terminal FLAG tag and a C-terminal cMyc tag. The resulting amino-acid sequence is:

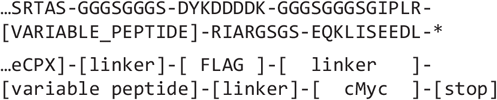

Individual peptides analyzed as shown in **Supplementary Figure 1** were cloned using Gibson assembly (Gibson *et al*., 2009) to replace the variable peptide sequence described above.

### Protein expression and purification

#### Expression of LC3B proteins

LC3B proteins were expressed in *E. coli* Rosetta2(DE3) cells (Novagen) grown at 37 °C with aeration via shaking in 2×YT medium supplemented with ampicillin (100 µg/mL) and D-(+)-biotin (0.05 mM). When cultures reached OD_600_ ≈ 0.8, protein expression was induced by addition of IPTG (1 mM), and cells were shifted to 25 °C and incubated overnight with aeration before harvesting by centrifugation (5000×g, 15 minutes, 4 °C). Cell pellets were flash-frozen and stored at -80 °C until purification. The GST-3C-BLM^554-571^-GS-LC3B fusion protein was expressed as above using *E. coli* BL21(DE3) cells.

#### Expression of peptides

Peptide constructs were expressed in *E. coli* BL21(DE3) cells (New England Biolabs) cultured with aeration at 37 °C in 20 mL Luria Broth (LB) supplemented with ampicillin (100 µg/mL) and D-(+)-biotin (0.05 mM). Expression was induced at an OD_600_ ≈ 0.8 using IPTG (1 mM) and, after 4-5 hours of expression at 37 °C, the cells were harvested and frozen as above.

#### Purification of LC3B proteins

Cell pellets were resuspended in buffer NB (20 mM Tris-HCl pH 8.0, 500 mM NaCl, 5 mM imidazole, 0.5 mM TCEP) supplemented with 0.2 mM phenylmethylsulfonyl fluoride (PMSF). Cells were lysed by 20 passes through a Dounce homogenizer followed by sonication on ice (4 × 2.5min cycles, each alternating 20 sec at 30% amplitude, 10 sec off ). The lysate was clarified by centrifugation (15,000×g, 15 min, 4 °C) and ultracentrifugation in a 60Ti rotor (50,000 rpm, 1 hr). The supernatant was loaded onto a 5 mL Bio-Scale Mini Nuvia IMAC Ni-charged cartridge (Bio-Rad 780-0811) equilibrated in buffer NB and eluted with a 0-100% linear gradient of buffer EB (20 mM Tris-HCl pH 7.5, 500 mM NaCl, 300 mM imidazole). Fractions containing LC3B, as identified by SDS-PAGE, were pooled, concentrated (3 kDa cutoff spin concentrator; Millipore), and further purified by size-exclusion chromatography on a Superdex 75 16/600 (Cytiva) column equilibrated with buffer GFB (20 mM Tris-HCl pH 7.5, 150 mM NaCl, 0.5 mM TCEP, 10 % glycerol). Purified protein was concentrated to ∼1.3 mM, flash-frozen in liquid nitrogen, and stored at -80 °C.

Cell pellets bearing BLM^554-571^-GS-LC3B were resuspended in buffer GLB (20 mM Na_2_HPO_4_, 1.8 mM KH_2_PO_4_, 140 mM NaCl, 2.7 mM KCl, 0.2 mM PMSF; pH 7.3) and lysed and clarified as above. Supernatant was then loaded onto a GSTPrep FF 16/10 10 mL column (Cytiva) pre-equilibrated with buffer GLB lacking PMSF, washed, and eluted with buffer GEB (50 mM Tris-HCl pH 8.0, 150 mM NaCl, 30 mM reduced glutathione). Fractions were analyzed by SDS-PAGE, and those containing the correct molecular-weight species were pooled. Cleavage was performed overnight at 4 °C using a 1:100 (w/w) ratio of 3C protease (gift of Dr. Andrew V. Grassetti) to substrate. The cleavage product was then purified via size-exclusion chromatography as above, concentrated to ∼1.4 mM, flash-frozen in liquid nitrogen, and stored at -80 °C in 50 μL aliquots.

#### Purification of peptides for BLI assays

Cell pellets were thawed on ice and lysed using 4 mL B-PER reagent (Thermo Fisher) per gram of cells, supplemented with 0.2 mM PMSF. Lysed cells (∼1 mL) were rotated at 25 °C for 15 min before pelleting by centrifugation at 15,000 rpm for 15 min. Supernatant was added to 250 μL Ni-Sepharose High Performance resin (Cytiva) pre-equilibrated with buffer NB in a gravity flow column (Bio-Rad). Resin was washed three times with 1 mL buffer NB before elution with 2 mL of buffer EB. Purified products were flash-frozen and stored at -80 °C.

### FACS sample preparation

FACS samples were prepared as previously described (Hwang *et al*., 2022) with slight modifications detailed below.

#### Bacterial display sample preparation

Briefly, MC1061 *E. coli* cells (Casadaban and Cohen, 1980) were transformed either with a control plasmid by chemical transformation or with the human peptidome library by electroporation. Transformed cells were cultured overnight with aeration at 37 °C in LB (5 mL) supplemented with chloramphenicol (25 µg/mL) and glucose (0.2% w/v). The following morning, ∼1×10^7^ cells per sample to be analyzed (as estimated using OD_600_, assuming 5×10^8^ cells/ mL/OD_600_) were pelleted (3,000×g, 5 min), and resuspended in Terrific Broth supplemented with chloramphenicol (25 µg/mL) to an OD_600_ ≈ 0.1. Cultures were grown with aeration at 37 °C until reaching an OD_600_ ≈ 0.5, at which point peptide expression was induced with 0.04% (w/v) arabinose for 1.5 hr at 37 °C with aeration. Following induction, cells (∼1×10^7^ per sample) were centrifuged (3,000×g, 5 min), washed once with phosphate-buffered saline (PBS; 10 mM Na_2_HPO_4_, 1.8 mM KH_2_PO_4_, 2.7 mM KCl, 138 mM NaCl; pH 7.4) containing 0.1% (w/v) bovine serum albumin (BSA), and resuspended in PBS + BSA (0.1% w/v) to a final cell density of 4×10^8^ cells/mL.

An anti-FLAG antibody conjugated to allophycocyanin (anti-FLAG-APC; PerkinElmer) was diluted 1:100 in PBS + BSA (0.1%), and 30 µL of solution was added per 1×10^7^ cells (25 µL/sample) for each sample to be analyzed. Sample was rotated at 4 °C for 30 min in foil-wrapped 1.5 mL tubes, pelleted (3,000×g, 5 min) and washed with 500 µL PBS + BSA (0.1 %) to remove unbound antibody, and resuspended in 25 µL PBS per sample to a final cell density of ∼4×10^8^ cells/mL.

#### LC3B sample preparation

Biotinylated BAP-6×His-LC3B proteins were bound to streptavidin-conjugated-phycoerythrin (SA-PE, Thermo Fisher Scientific) at a 4.2:1 molar ratio in PBS+BSA (1.0 %) containing 4 mM dithiothreitol (DTT). Complexes were incubated in the dark with rotation for 15 min at 4 °C and prepared at 2× the desired final concentration. For all sorting experiments, LC3B-SA-PE was prepared to yield a final concentration of 1.68 µM monomeric LC3B (0.42 µM tetramerized LC3B•SA-PE).

#### FACS sample preparation

To prepare final samples, 25 µL of cells (1×10^7^) in PBS was mixed 1:1 with 25 µL of pre-tetramerized LC3B (2×stock concentration) for a final volume of 50 µL, and incubated with rotation in the dark at 4 °C for 1 hr. Note that for the initial library, 200 µL of cells (∼1×10^8^) were mixed 1:1 and treated as above. Samples were transferred to a 0.22 µm 96-well Multi-Screen HTS GV sterile filtration plate (Millipore) pre-washed with 500 µL PBS + BSA (0.5%) per well. Cell suspensions were vacuum filtered, washed with 200 µL PBS + BSA (0.1%) and resuspended in 400 µL PBS + BSA (0.1%) for control experiments shown in Supplementary Figure 1 or 4.2 mL PBS + BSA (0.1%) for library samples prior to FACS analysis or sorting.

### Bacterial display sorting

Bacterial display libraries were analyzed and sorted on the BD FACSCanto II (analysis) and BD FACSAria 4 (sorting) instruments. To establish gating parameters for enrichment of LC3B-binding peptides, two positive controls (FYCO1^1264-1299^ and ATG4B^358-393^) and a negative control (lacking the variable peptide segment) were analyzed. The selection gate was set to maximize recovery of positive cells while maintaining ≤ 0.4 % background from the negative population. Following fluorescence compensation to correct for spectral overlap between channels, the gate was held constant throughout all positive enrichment sorts.

In the initial sort, ∼8.6×10^7^ cells were sorted – sufficient to oversample the input library by ∼200-fold. Sorted cells were collected into Super Optimal broth with Catabolite repression (SOC) and incubated for 1 hr at 37 °C with rotation before being transferred to 8 mL LB medium containing chloramphenicol (25 μg/mL) and glucose (0.2 % w/v). Cultures were grown at 37 °C with aeration until OD_600_ ≈ 0.4–0.8. Plasmid DNA was purified from these cells using a Monarch Miniprep Kit (New England Biolabs).

For each subsequent round of sorting, *E. coli* MC1061 cells were transformed via electroporation with 50 ng of the purified plasmid library DNA and analyzed as above, with sufficient cells collected to oversample the binding pool by ∼100-fold. For the second round (sort 2), which served as a counter-selection against nonspecific peptide binding to the assay components, D-(+)-biotin was incubated with SA-PE at a 4.2:1 molar ratio as described above, and a gate was selected that minimized the capture of positive-control peptides (**Supplementary Figure 1)**.

LC3B LDS* samples were prepared in a similar manner as wild-type LC3B. To enable direct comparison, the binding pool from sort 5 was split and sorted in parallel on the same day against wild-type LC3B (sort 6) and LC3B LDS* (LDS* 1). Both binding and non-binding populations from each experiment were collected for sequencing. Two additional LC3B LDS* enrichment sorting rounds were then performed to further enrich for LDS*-binding peptides, and both the binding and non-binding populations were collected and miniprepped to isolate the plasmid DNA.

### Next-generation sequencing (NGS) sample preparation

Miniprepped plasmid libraries from each round of sorting were assigned a unique 6-nucleotide index. The variable region of each library was PCR-amplified using a forward primer (ol_JD806) bearing the 5′ Illumina adaptor sequence (AATGATACGGCGACCACCGAGATCTACAC) and a sequence (GTGGCTCGGGAATTCCGCTGCGC) designed to anneal immediately 5′ of the variable peptide region of our bacterial display vector, and a reverse primer (ol_JD807-813) designed to anneal immediately 3′ of the variable peptide region that also contained one of the fourteen 6-nucleotide index sequences and a 3′ Illumina adaptor sequence. The amplicon preparation scheme and all primers are listed in **Supplementary File 1**.

Eleven cycles of amplification were performed using Phusion polymerase (NEB) with 1.25 µM primers under the following thermocycling conditions: initial denaturation at 98 °C for 45 s; denaturation at 98 °C for 15 s; annealing at 68 °C for 30 s; and extension at 72 °C for 60 s, followed by a final extension at 72 °C for 5 min. To minimize heteroduplex formation, two additional cycles of reconditioning PCR were performed with 1 µM primers using the same thermocycling conditions. PCR products were purified using a double-sided AMPure XP (Beckman Coulter) bead size selection (0.85X/0.5X) to exclude large amplicons and small adapter or primer dimers, retaining fragments within the desired library size range for Illumina sequencing. Samples were eluted in 11 μL of buffer NGS (10 mM Tris-HCl, pH 8.0).

DNA amplicon library purity, size, and quantity were assessed using a Bioanalyzer (Agilent Technologies). Indexed samples were pooled to generate a multiplexed library bearing samples from all wild-type LC3B and LC3B LDS* sorts, which was submitted to the MIT BioMicro Center for sequencing on a NextSeq500 (Illumina) platform using 150 bp paired-end reads.

### NGS data processing and analysis

Demultiplexed sequencing data were merged with BBMerge (Bushnell *et al*., 2017) using an average quality score threshold of 20. Merged reads were used to quantify the number of clonal cells (*i*.*e*., cells displaying the same peptide) recovered across consecutive sorts. The frequency of each clone *i* collected in sort *x*, denoted as *f*_*i,x*_, was calculated as (*c*_*i,x*_ */ N*_*x*_) where *c*_*i,x*_ is the number of sequencing reads mapped to sequence *i* in sort *x* and *N*_*x*_ is the total number of sequencing reads mapped in sort *x, i*.*e*., *N*_*x*_ *= Σ*_*i*_ *c*_*i,x*_. The enrichment ratio (ER) for each clone across the sorting trajectory was then calculated as ER_*i,x*_ = log_2_(*f*_*i,x*_ / *f*_*i,input*_) where *f*_*i,input*_ represents the frequency of the same sequence in the naïve input library (Rubin *et al*., 2017).

The T7-pep library contains many apparent point mutations and frameshifts. To group related variants, sequences were clustered using ALFAT-Clust (Chiu and Ong, 2022) with default parameters. Within each cluster, the variant that persisted to the latest sort was selected as the representative sequence; when multiple variants persisted, the sequence with the highest read count in that sort was chosen. Clones with fewer than 10 reads in the input library were excluded from all analyses. Sequences were translated using Biopython (Cock *et al*., 2009) and mapped to the human proteome using NCBI BLASTP (Camacho *et al*., 2009).

All read count data for each sequence in each sort are available at the NCBI Sequence Read Archive (SRA) under BioProject ID PRJNA1276872. **Supplementary File 2** contains the pre-collapsed data, and **Supplementary File 3** contains the post-collapsed data used in this study, including amino acid sequences, raw counts, ER values, BLAST results, and *z*-scores. **Supplementary File 4** is filtered for peptides that enriched through sort 6 and includes AlphaFold2 (Jumper *et al*., 2021) pLDDT scores estimating structural disorder within each peptide region.

### LC3B and LC3B LDS* scoring metrics

Following NGS data processing described above, 487,021 unique peptide sequences were identified in the input library, with 12,158 remaining in the terminal LC3B sort 6. Among these, clones were placed in the HC-set if the mean *z*-score across sorts 4-6 was ≥1.70. The *z*-score for clone *i* in sort *x* was defined as: *z*_*i,x*_ = [(ER_*i,x*_ - μ_*x*_) / σ_*x*_], where ER_*i,x*_ is the enrichment ratio of peptide *i* in sort *x* and μ_*x*_ and σ_*x*_ are the mean and standard deviation ER values for all peptides in that sort. The threshold of 1.70 was chosen based on BLI affinity measurements to minimize the false positives (**Figure 2C**). *Z*-scores for all clones that reached sort 6 are provided (**Supplementary File 4**).

To evaluate peptide behavior in the LC3B LDS* sorts, three complementary metrics were applied. First, enrichment trajectories were examined to identify clones that had enriched through the canonical LC3B sort 5, and that continued to enrich through the three LDS* sorts despite disruption of the canonical LIR-docking site. Second, the enrichment ratio was calculated using sort 5 as the input library to compare peptide enrichment between the first LDS* sort (ER LDS*_i,1_ = freq(LDS*_i,1_) / freq(sort 5)) and the canonical sort 6 (ER_i,6_ = freq(sort 6) / freq(sort 5)). Promising clones were those that were enriched in the LDS* condition and depleted in the canonical condition (ER LDS*_i,1_/ ER_i,6_ > 1), consistent with binding in the absence of the wild-type LDS pocket. Finally, the ratio of clone frequencies in the binding versus non-binding gates (B/NB) was calculated for each LDS* sort and for the canonical LC3B sort 6. Clones showing progressive increase in the B/NB ratio across the LDS* sorts were considered selectively enriched for binding to LC3B LDS*. Top-scoring clones across these metrics were manually selected and tested for binding to LC3B and LC3B LDS* by BLI, as detailed below. Peptides that bound LC3B LDS* scored highly across all three metrics. All LDS* enrichment data and associated scoring metrics are provided (**Supplementary File 5**).

### Identification of reported interaction partners of LC3B

To assemble a set of high-confidence LC3B-binding peptides and their corresponding human proteins, the LIRcentral database (Chatzichristofi *et al*., 2023) was used to extract entries (n=115) experimentally verified to bind one of the hAtg8 proteins. Of these, 27 were reported to bind LC3B, with 10 such peptides reported to bind with K_D_ ≤ 50 µM being present in our input library. The six peptides from this subset that appeared in our HC-set are indicated in the “LIR row” of **Figure 2A**.

In parallel, human proteins reported to co-immunoprecipitate with LC3B (n=550) were retrieved from the BioGrid database (Stark *et al*., 2006), and peptides related to those proteins that were present in our HC-set are annotated in the “IP row” in **Figure 2A**. To assess whether candidates in the HC-set may have biologically relevant interactions with LC3B, Gene Ontology (GO) terms for proteins corresponding to each peptide present in sort 6 were extracted from UniProt (The UniProt Consortium, 2025) and cross-referenced with GO terms associated with LC3B. Those proteins sharing a GO term with LC3B are noted in the GO row of **Figure 2A**, with the GO terms shown in **Supplementary Figure 4**.

### Biolayer interferometry (BLI)

BLI experiments were performed on an Octet Red96 instrument (ForteBio). Purified, biotinylated SUMO-fused peptides were immobilized on Octet SA Biosensors (Sartorius) and loaded until a response level of at least 0.3 nm was reached. The ligand-loaded tips were then incubated with increasing concentrations of target protein (*e*.*g*., LC3B) in buffer BLI (25 mM Tris, pH 7.5, 150 mM NaCl, 0.5 mM TCEP, 0.05 % Tween-20, 0.5 mg/mL BSA) in a 200 μL reaction conducted in a 96-well flat-bottom polypropylene microplate (Greiner, #655209). Each measurement was repeated 2 – 4 times at 25 °C at an agitation speed of 1000 RPM. Equilibrium dissociation constants (K_D_) for the measured interactions were determined by plotting the equilibrated signal, after subtracting the negative control (biotinylated SUMO-only), as a function of protein concentration. The data were fit by nonlinear regression to a one-site binding model, according to the equation:

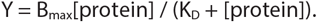

### Structure determination by X-ray crystallography

Crystals of the purified BLM^554-571^-GS-LC3B fusion protein were grown in hanging drops containing buffer BXC (0.1 M HEPES, pH 7.3, PEG-3350 (30 % w/v), and 0.32 M MgCl_2_), by mixing 1 μL of protein with an equal volume of reservoir solution; crystals appeared within 7 days. X-ray diffraction data were collected on a Rigaku Micromax-007 rotating anode equipped with Osmic VariMax-HF mirrors and a Rigaku Saturn 944 detector. Diffraction data were processed with the XDS suite (Kabsch, 2010).

The structure was solved by molecular replacement in Phaser (McCoy *et al*., 2007) using LC3B (pdb 3vtu) (Rogov *et al*., 2013) as the search model. The molecular replacement solution was refined in PHENIX (Liebschner *et al*., 2019) with manual fitting in Coot (Emsley *et al*., 2010), and further geometry optimization in Rosetta (DiMaio *et al*., 2013). The model was refined to R_work_/ R_free_=0.220/0.263. The X-ray data collection and refinement statistics are summarized in **Supplementary Table 4**, and the coordinates have been deposited in the Protein Data Bank (pdb 9p3e). Ensemble refinement as implemented in PHENIX (Burnley *et al*., 2012) was carried out using pTLS values (fraction of atoms included in TLS fitting) ranging from 1.0 to 0.5, with 0.6 yielding the lowest R_free_ value.

### Sequence logo-based analysis

We used pLogo (O’Shea *et al*., 2013) to visualize the enrichment of residues in peptide binders, compared to the input library background, and designed a series of consensus-sequence peptides based on this analysis. pLogo estimates the statistical significance of residue enrichments using foreground and background distributions. To define the background, we identified input-library sequences matching the LIR motif and included 15 residues on either side of the motif. For peptides that lacked 15 flanking residues around the LIR, we appended the N- or C-terminal flanking sequence that was present in the experimental display construct. To define a set of strongly enriching binders for the foreground, peptides that reached sort 6 were clustered (k=10), based on their enrichment profiles, using Clust (Abu-Jamous and Kelly, 2018). The sequences in the highest-enriching cluster were used as the foreground, using the LIR plus 15 flanking residues on either side, as above. The foreground and background sequences were aligned on the LIR and used as inputs to pLogo. We repeated this analysis four times, for the canonical [FWY]_0_-X_1_-X_2_-[LVI]_3_ LIR motif and for LIR motifs with position X_0_ restricted to F, W, or Y, generating the images in **Supplementary Figure 5**. For consensus peptide design, the residue with the highest log-odds of the binomial probability at each position of the corresponding 34-residue logo was selected, generating peptides pCONSLIR, pCONS_F_, pCONS_W_, and pCONS_Y_, which are listed in **Supplementary Table 3**.

### LIR sequence sets for PSSM and random-forest model building

To build and assess models for classifying LC3B-binding and non-binding peptides, five different sequence sets were defined, each matching the 7-residue motif X_-3_X_-2_X_-1_[FWY]_0_X_1_X_2_[FWYILV]_3_: *screen_binder* and *screen_nonbinder* from our screening data, *test_binder* and *test_nonbinder* sequences from LIRcentral (test sets) (Chatzichristofi *et al*., 2023), and *iLIR_binder* sequences from iLIR (Kalvari *et al*., 2014). To define the *iLIR_binder* set, LIR-containing sequences were downloaded from the iLIR webserver. Because the iLIR motifs are only 6 residues long, we extended the 6-mers to 7-mers using full-length sequences from the UniProt reference proteome (The UniProt Consortium, 2025). To define the *test_binder* set, we extracted 76 motif-matching 7-mer sequences annotated to bind LC3B in LIRcentral and that were not present in the *iLIR_binder* set, then added three peptides from our screen that we measured to bind LC3B with K_D_ ≤ 14 µM using BLI (**Supplementary Table 1**). To define the *test_nonbinder* set, we used 49 motif-matching 7-mers reported by LIRcentral to not bind LC3B (also absent from the *iLIR_binder* set), then added seven peptides from our screen with no detectable binding to LC3B by BLI (**Supplementary Table 2**). In total, the test set contained 79 binder peptides and 56 non-binder peptides.

To define the *screen_binder* and *screen_nonbinder* sets, we selected input-library sequences bearing exactly one LIR motif (as above), appended “PLR” to the N-terminus, which was present in the experimental display construct, extracted 7-mers centered on the LIR motif, and removed any sequences also present in the *iLIR_binder, test_binder*, or *test_nonbinder* sets. Remaining 7-mers in the HC-set formed the *screen_binders* set; those that were depleted in sorts 1 and 3 and then dropped out of the screen during any subsequent sort formed the *screen_nonbinders* set. Note that any duplicate 7-mers found within or between the *screen_binder* and *screen_nonbinder* sets were removed from both sets. The final sequence set from the screening data consisted of 151 *screen_binder* sequences and 2,844 *screen_nonbinder* sequences. For modelling, we additionally used ranges of *z*-scores to define *screen_binder* subsets: *screen_binder_all* (*z*-score ≥ 1.7, 151 sequences), *screen_binder_low* (1.7 ≤ z-score < 2.3, 101 sequences), and *screen_binder_high* (z-score ≥ 2.3, 50 sequences). Logos for the screening sets are shown in **Supplementary Figure 6C**.

For the model building and evaluation, screening data were repeatedly (100×) split into train/test partitions. Each iteration selected 7-mers at random without replacement to form test sets, with 15 from the *screen_binder_high* test subset, and 30 from the *screen_binder_low* test subset; we designated the combined 45 sequences as the “all” *screen_binder* test set. An equal number of nonbinder sequences from the *screen_nonbinder* set was selected to complete the test sets. For models built from the screening data, the sequences that were not selected for the test sets were used to train models or build PSSMs. Thus, for each train-test iteration, three completely non-overlapping test and training sets were defined, which were used to train and evaluate models.

### PSSM classifier model construction

PSSM classifier models were generated using a 7-residue window spanning the three N-terminal flanking residues and the core LIR motif (*i*.*e*., X_-3_X_-2_X_-1_[FWY]_0_X_1_X_2_[FWYILV]_3_). Each model used one of the binder sets (defined above), as the foreground and the amino-acid frequencies from the *Homo sapiens* reference proteome (UP000005640) as the background distribution. Position-specific scores were calculated as log-odds values:

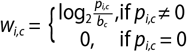

where *w*_*i,c*_ is the position-specific score for residue *c* at position *i, p*_*i,c*_ is the frequency of residue *c* at position *i* in the set of foreground sequences, and *b*_*c*_ is the background frequency of residue *c* in the proteome. The total PSSM score for a sequence was the sum of position-specific values across the 7-residue window. An analogous model (iLIR_27_) was generated using the 27 binders that formed the basis of the published iLIR model (Kalvari *et al*., 2014) as the foreground sequences.

### Random-forest classifier model construction

Random-forest and balanced random-forest classifiers were trained using *RandomForestClassifier* from scikit-learn (Pedregosa *et al*., 2011) and *BalancedRandomForestClassifier* from imbalanced-learn (Lemaître *et al*., 2017), where we noted that unlike the PSSM model from above, these models have the potential to capture higher-order effects, including coupling between sites. The training set consisted of the 2,844 nonbinder and 151 binder sequences defined above. We leveraged the PSSM matrix detailed above to encode each sequence. A PSSM-encoded sequence (X) is generated by replacing the one-letter amino acid code in a sequence with its corresponding PSSM score in the matrix. Mathematically, the *i*^th^ encoded position of a sequence (X_*i*_) is:

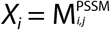

where *i* denotes the *i*^th^ position in an encoded peptide sequence, *j* represents the column in the PSSM matrix that corresponds to the amino acid appearing at *i*^th^ position in a sequence. To handle data imbalances between screening binders and nonbinders, a downsampling technique was used to randomly select nonbinders to match the total number of binders. Cross-validation (5-fold) was applied to select the best hyperparameters from the parameter space for random forest (*n_estimators: 4-200, criterion: [gini, entropy], ccp_alpha: 0*.*004-0*.*014*) and balanced random forest models (*n_estimators: 4-200, criterion: [gini, entropy], ccp_alpha: 0*.*004-0*.*014, sampling_strategy: all, replacement: True, bootstrap: False*). The best model was selected based on the mean ROC-AUC and retrained with all training data.

## ACKNOWLEDGEMENTS

We thank A. Ghanbarpour for help with crystallographic structure refinement. We thank L. Kinman, D. Cui, and B. Powell for helpful discussions. Research reported in this publication was supported by the National Institutes of Health under Award Numbers R35GM149227 (AEK), R01GM144542 (JHD), R00AG050749 (JHD) and the Smith Family Odyssey Award (JHD).

## AUTHOR CONTRIBUTIONS

J.E.K., J.H.D., and A.E.K. designed research; J.E.K. performed screening, data analysis, and biochemical experiments, and contributed new reagents and analytic tools; C.L., J.E.K., and J.C.H. developed and tested models; D.L. collected and refined crystallography data; J.E.K., J.H.D., and A.E.K. wrote the paper.

## COMPETING INTERESTS

The authors declare no competing interests.

## DATA AVAILABILITY

Datasets generated during this study have been deposited in the National Center for Biotechnology Information Sequence Read Archive (NCBI SRA). Raw reads can be found under BioProject ID PRJNA1276872. The structure of LC3B bound to BLM^554-571^ is deposited in the Protein Data Bank with accession number 9p3e. Supplementary Files 1-5 are available at Zenodo (<u>10.5281/zenodo.17943925</u>). Python processing scripts are available upon request.

*Binding determinants of human LC3B*

## Supplementary Information

## SUPPLEMENTARY TABLES

**Supplementary Table 1.**
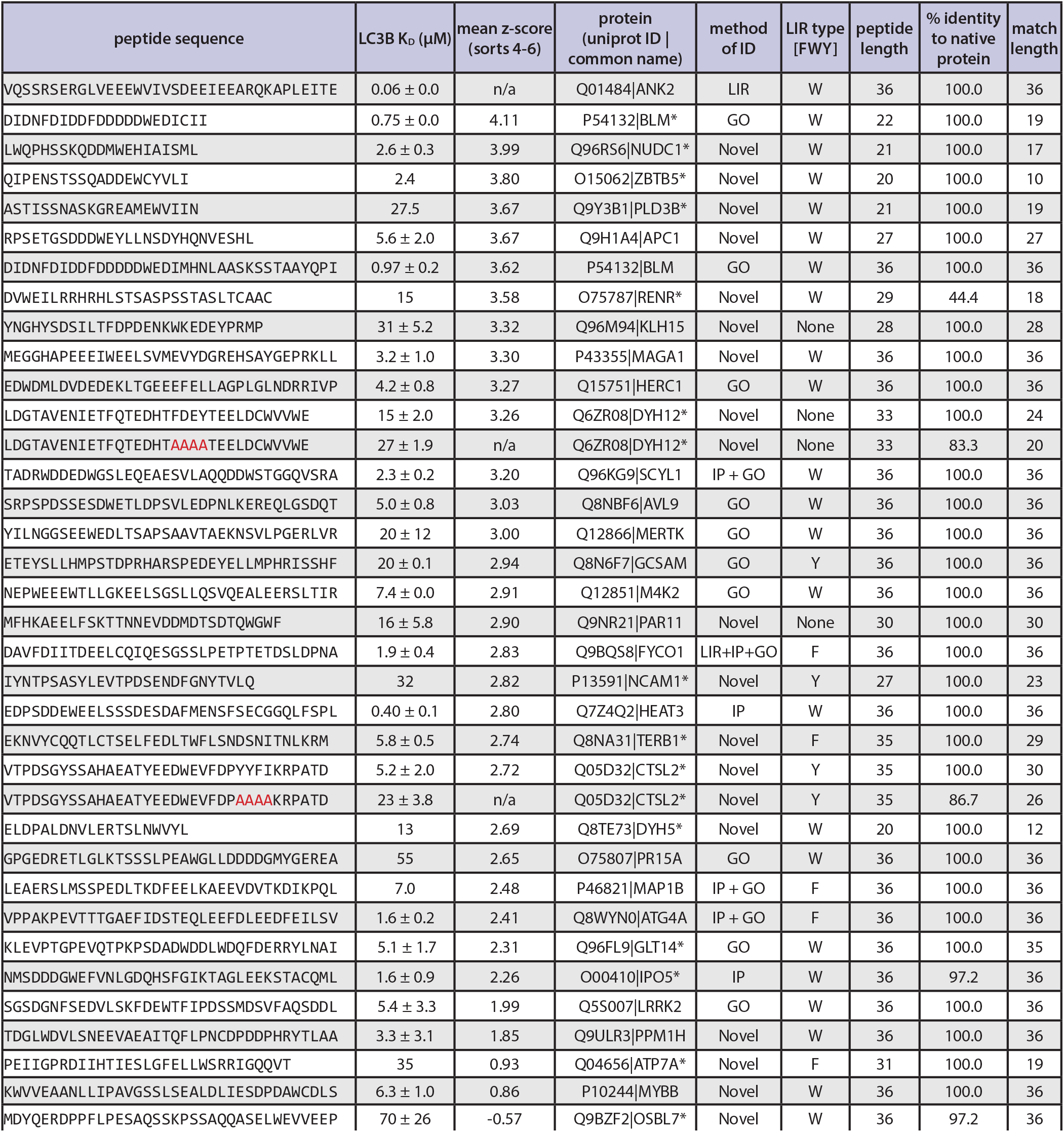
LC3B binding affinities determined using bio-layer interferometry (BLI). K_D_ values report the average and standard deviation of n ≥ 2 replicates; if no standard deviation is noted, only one replicate was measured for the given peptide. For each peptide, a BLAST search was performed to determine the corresponding protein, with the percentage sequence identity calculated over the length of the sequence match listed in the final two columns. Peptides bearing mutations or truncations are noted with *. Peptides with alanine substitutions in putative LIR motifs are highlighted in red.

**Supplementary Table 2.**
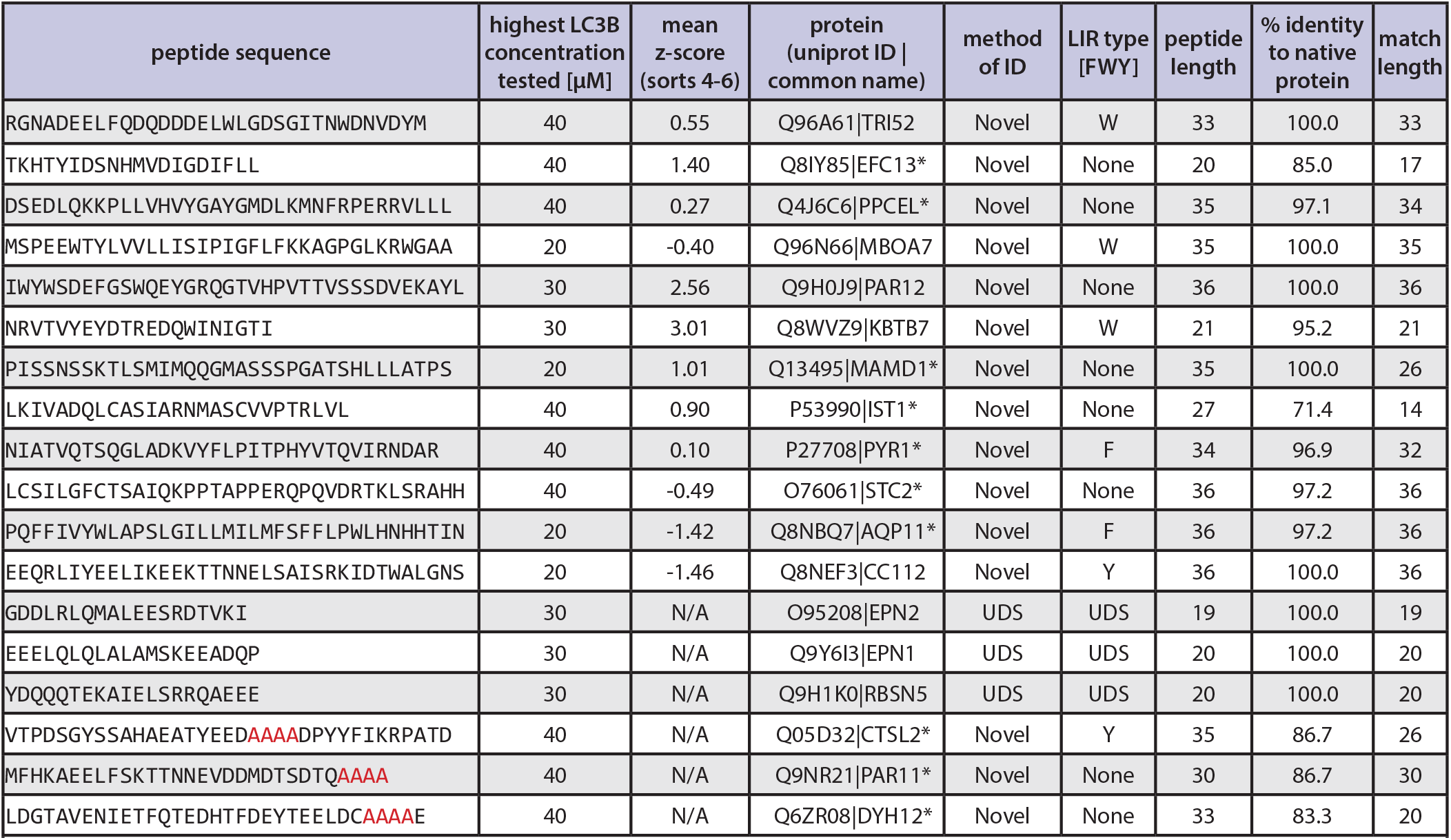
Peptides without observable binding to LC3B. Peptides were tested for binding to LC3B by BLI at the noted concentrations, and no binding was observed. Peptides predicted to bind at the “UDS” (Marshall *et al*. 2019) are marked with LIR type UDS. Peptides with alanine substitutions in putative LIR motifs are highlighted in red.

**Supplementary Table 3.**
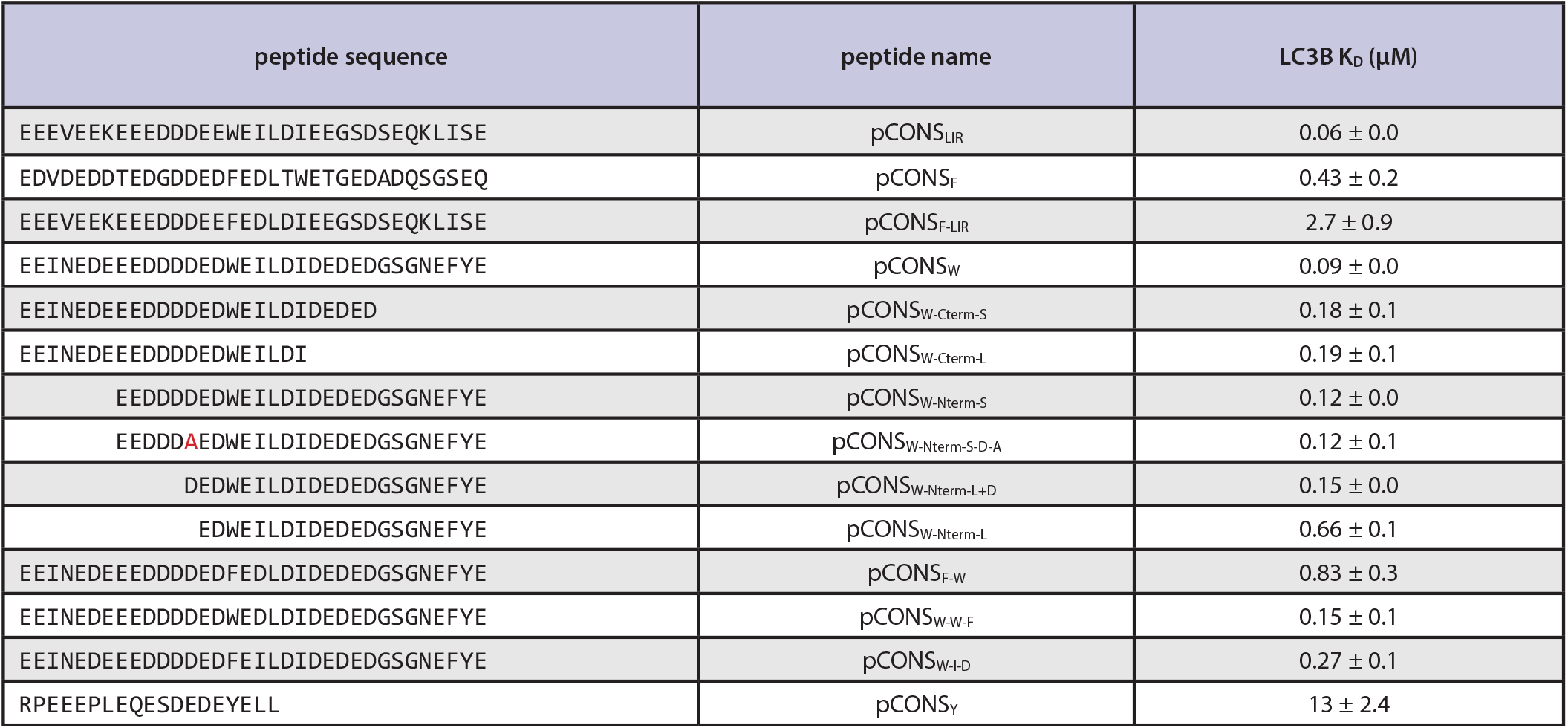
LC3B binding affinities of consensus peptides and related mutants. Binding affinity to LC3B measured for each pCONS peptide and various derivatives. K_D_ values are reported as the mean ± s.e.m. n ≥ 3 replicates.

**Supplementary Table 4.** Data collection and refinement statistics for the BLM-LC3B X-ray crystal structure (pdb 9p3e). Statistics for the highest-resolution shell are shown in brackets.

| Data collection and refinement statistics (molecular replacement) |  |
| --- | --- |
| Wavelength (Å) | 1.542 |
| Resolution range (Å) | 23.8 - 2.0 [2.1 - 2.0] |
| Space group | C 1 2 1 |
| Unit cell (Å, Å, Å; °, °, °) | 66.09 46.74 45.48; 90 102.2 90 |
| Total reflections | 49,019 [1,844] |
| Unique reflections | 9,065 [628] |
| Multiplicity | 5.4 [2.9] |
| Completeness (%) | 91.76 [72.79] |
| Mean I/sigma(I) | 10.91 [1.16] |
| Wilson B-factor (Å<sup>2</sup>) | 35.76 |
| R<sub>merge</sub> | 0.089 [0.874] |
| R<sub>meas</sub> | 0.098 [1.067] |
| R<sub>pim</sub> | 0.040 [0.597] |
| CC<sub>1/2</sub> | 0.998 [0.291] |
| CC* | 1 [0.672] |
| Reflections used in refinement | 9,062 [628] |
| Reflections used for R-free | 908 [61] |
| R<sub>work</sub> | 0.220 [0.376] |
| R<sub>free</sub> | 0.263 [0.404] |
| Number of non-hydrogen atoms | 1,183 |
| macromolecules | 1,117 |
| ligands | 1 |
| solvent | 65 |
| Protein residues | 135 |
| RMS<sub>bonds</sub> (Å) | 0.002 |
| RMS<sub>angles</sub> (°) | 0.54 |
| Ramachandran favored (%) | 97.74 |
| Ramachandran allowed (%) | 2.26 |
| Ramachandran outliers (%) | 0.00 |
| Rotamer outliers (%) | 2.34 |
| Clashscore | 0.45 |
| Average B-factor (Å<sup>2</sup>) | 51.65 |
| macromolecules (Å<sup>2</sup>) | 51.95 |
| ligands (Å<sup>2</sup>) | 51.52 |
| solvent (Å<sup>2</sup>) | 46.54 |

## SUPPLEMENTARY FIGURES

**Supplementary Figure 1.**
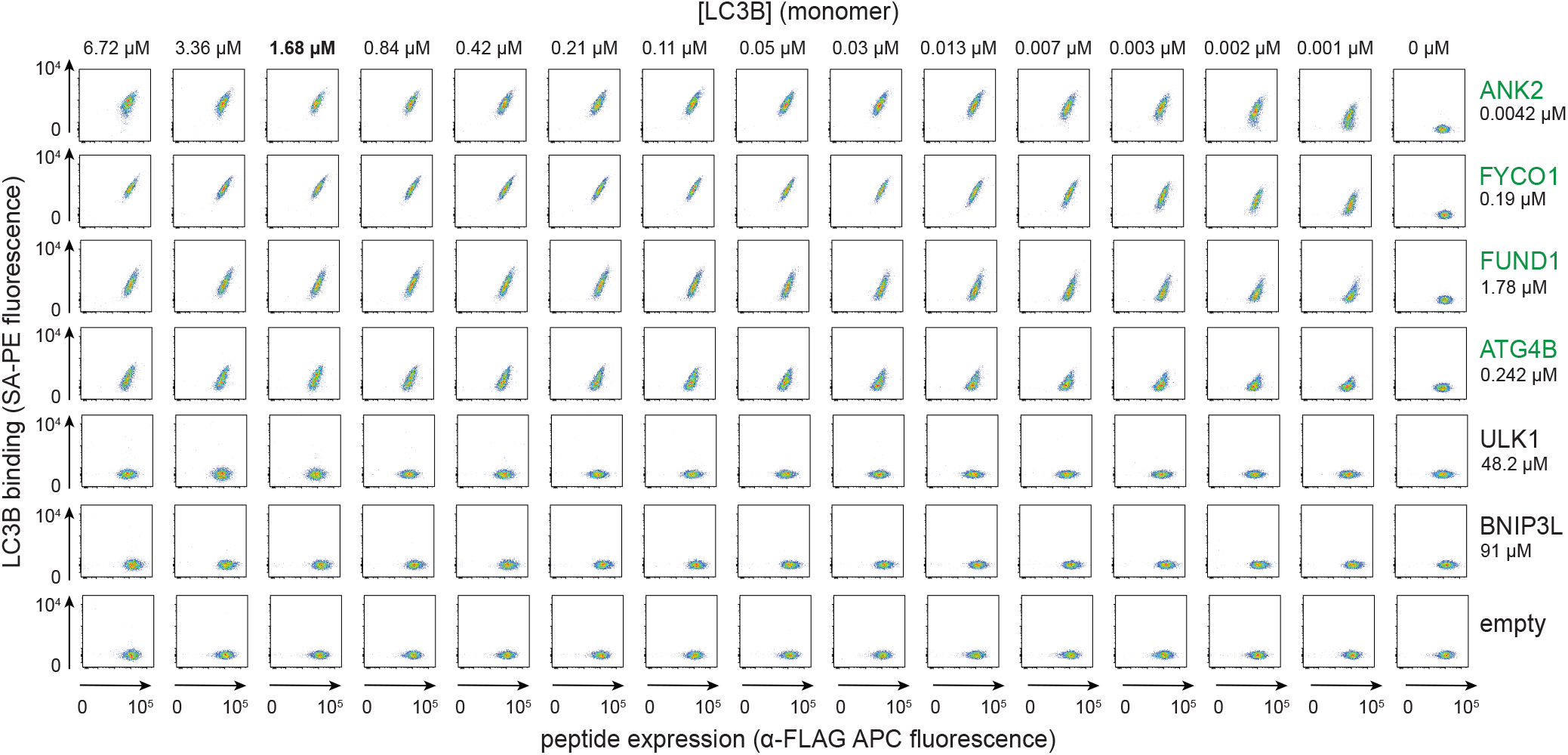
Dynamic range of well-studied LIR motifs binding to pre-tetramerized LC3B in bacterial display. Control peptides reported to bind LC3B: ANK2^1578-1613^ (VQSSRSERGLVEEEWVIVSDEEIEEARQKAPLEITE), FYCO1^1264-1299^ (GGQGANTDYRPPDDAVFDIITDEELCQIQESGSSLP), FUND1^2-37^ (ATRNPPPQDYESDDDSYEVLDLTEYARRHQWWNRVF), ATG4B^358-393^ (HLACPDVLNLSLDSSDVERLERFFDSEDEDFEILSL), ULK1^341-376^ (SRDSGGSKDSSCDTDDFVMVPAQFPGDLVAEAPSAK), BNIP3L^20-55^ (EENEQSLPPPAGLNSSWVELPMNSSNGNDNGNGKNG), and negative control (eCPX-FLAG with no peptide displayed) were assayed for binding to serially diluted pre-tetramerized LC3B using bacterial display. FACS profiles relating peptide expression and LC3B binding are plotted with points reflecting cell density, colored from blue to red to indicate increasing density. Peptides labeled green (right) exhibited concentration-dependent, saturable binding. Affinities listed below each protein name were obtained from papers referenced in LIRcentral (Chatzichristofi *et al*. 2023); note that the start and end positions of peptides assayed here may differ from those used in the cited studies. Based on these controls, a working concentration of 1.68 µM monomeric LC3B (0.42 µM when tetramerized by streptavidin) was selected for the proteome-wide screen and is indicated in bold font.

**Supplementary Figure 2.**
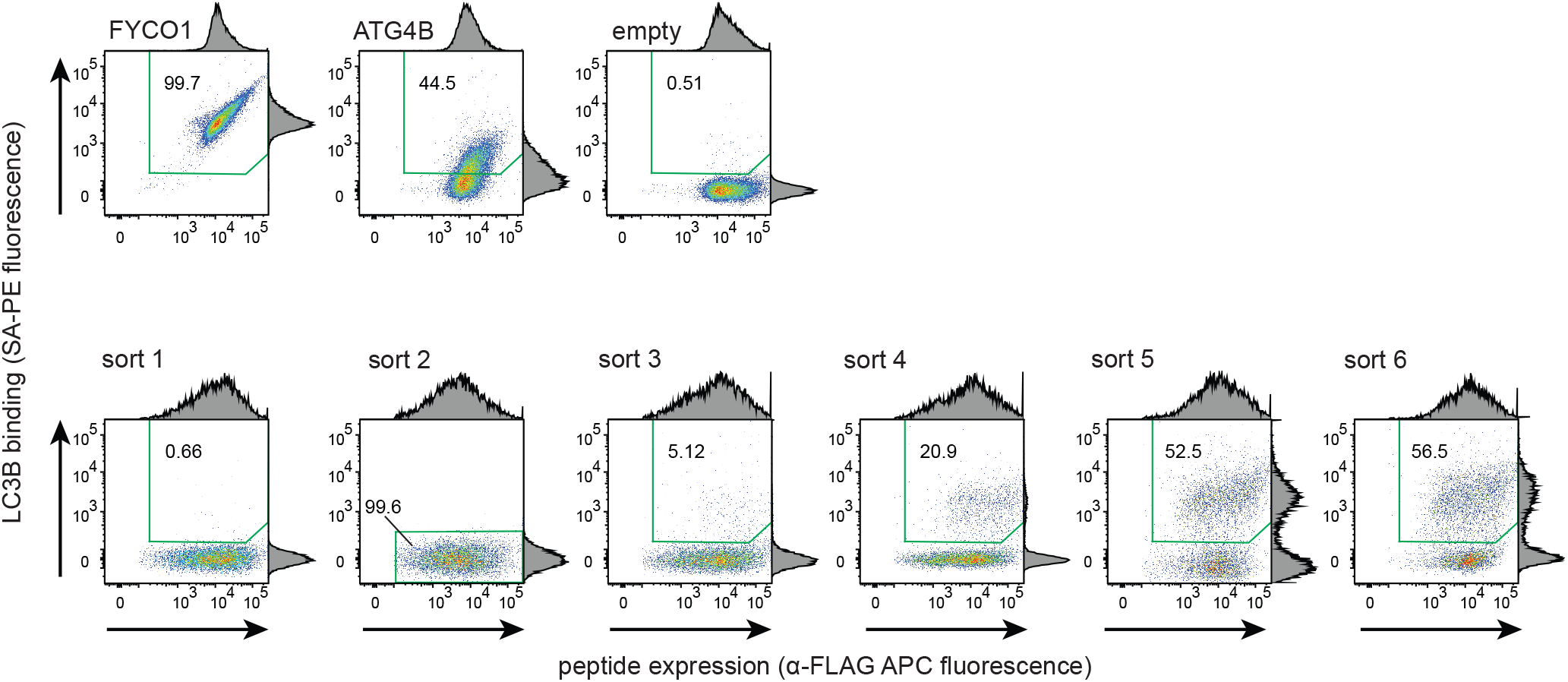
Bacterial display-based enrichment of peptides capable of binding pre-tetramerized LC3B. FACS profiles with marginal distributions relating peptide expression (y-axis) and LC3B binding (x-axis) are plotted with points reflecting cell density, colored from blue to red to indicate increasing density. The top row displays profiles measured during sort 1 for positive controls FYCO1^1264-1299^ and ATG4B^358-393^ and the negative control (eCPX-FLAG with no peptide displayed), labeled empty. The bottom row displays the results of sorting the input library for six rounds, selecting for binding to streptavidin-tetramerized LC3B. Note that sort 2 was designed to remove clones that bound to SA-PE in the absence of LC3B and uses a different selection gate (outlined in green) than the other five positive-selection sorts. The percentage of all analyzed cells captured in each gate is noted.

**Supplementary Figure 3.**
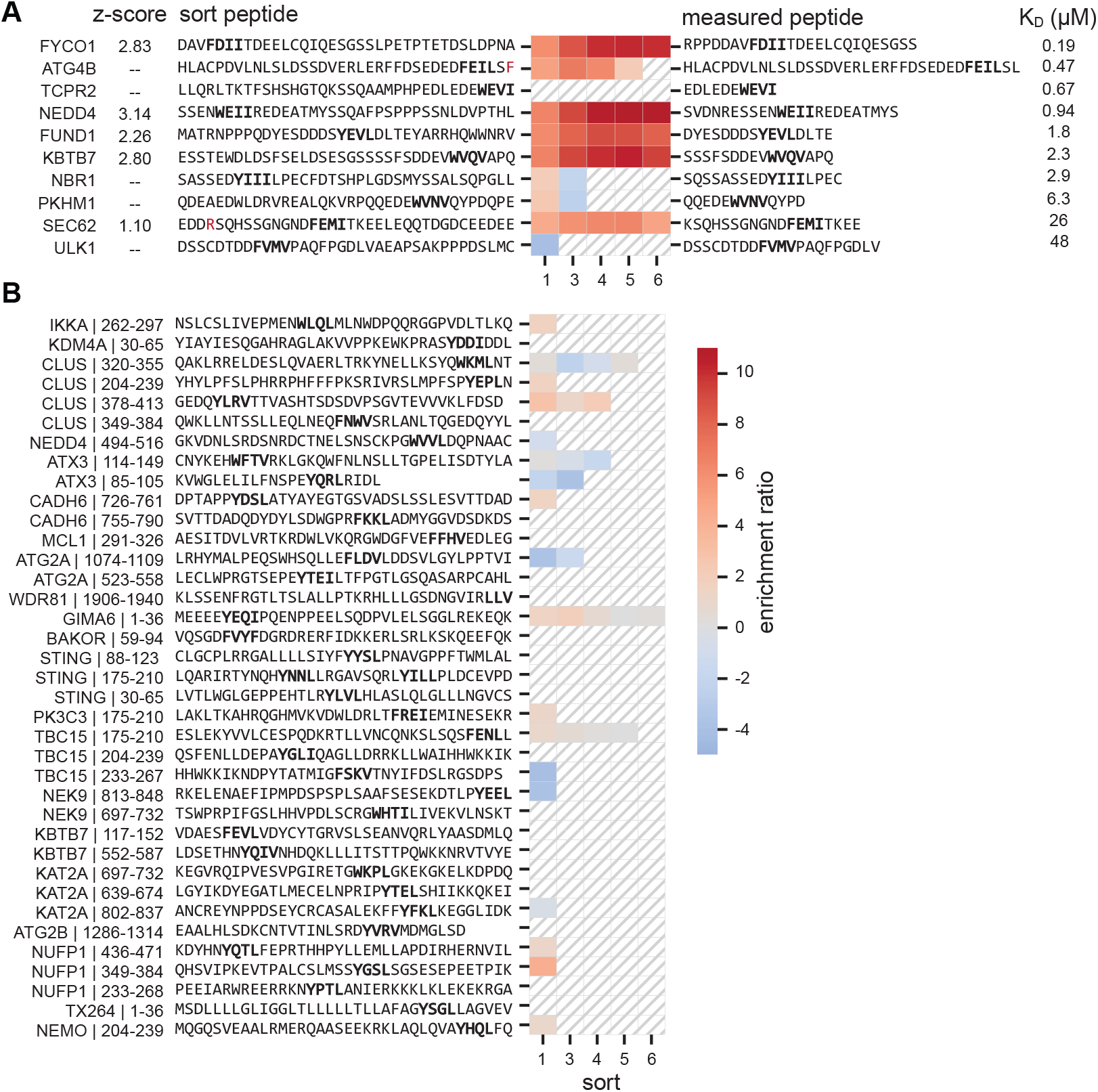
Sorting behavior of reported LC3B-binding peptides. Heatmap depicting the enrichment ratio of LIRcentral-annotated peptides in the input library (rows) observed in each round of sorting. Hashing indicates that the peptide was not identified in that sort. **(A)** High-affinity binders. The peptide identified from the sort is shown with the core LIR in bold and any mutations from the human sequence in red. The peptide tested for binding to LC3B is shown on the right. Previously measured affinities, and measurement techniques are as follows: FYCO1^1273-1297^ (0.19 µM; ITC) (Cheng *et al*., 2016), ATG4B^358-393^ (0.47 µM; BLI), TCPR2^1403-1411^ (0.67 µM; ITC) (Stadel *et al*., 2015), NEDD4^256-278^ (0.94 µM; FP) (Qiu *et al*., 2017), FUND1^10-25^ (1.8 µM; ITC) (Lv *et al*., 2017), KBTB7^658-674^ (2.3 µM; ITC) (Genau *et al*., 2015), NBR1^722-739^ (2.9 µM; ITC) (Rozenknop *et al*., 2011), PKHM1^629-642^ (6.3 µM; ITC) (Rogov *et al*., 2017), SEC62^352-370^ (26 µM; BLI) (Fumagalli *et al*., 2016), ULK1^349-369^ (48 µM; BLI) (Wirth *et al*., 2019). **(B)** Peptides annotated as non-binding in LIRcentral (Chatzichristofi *et al*., 2023), analyzed as in panel A.

**Supplementary Figure 4.**
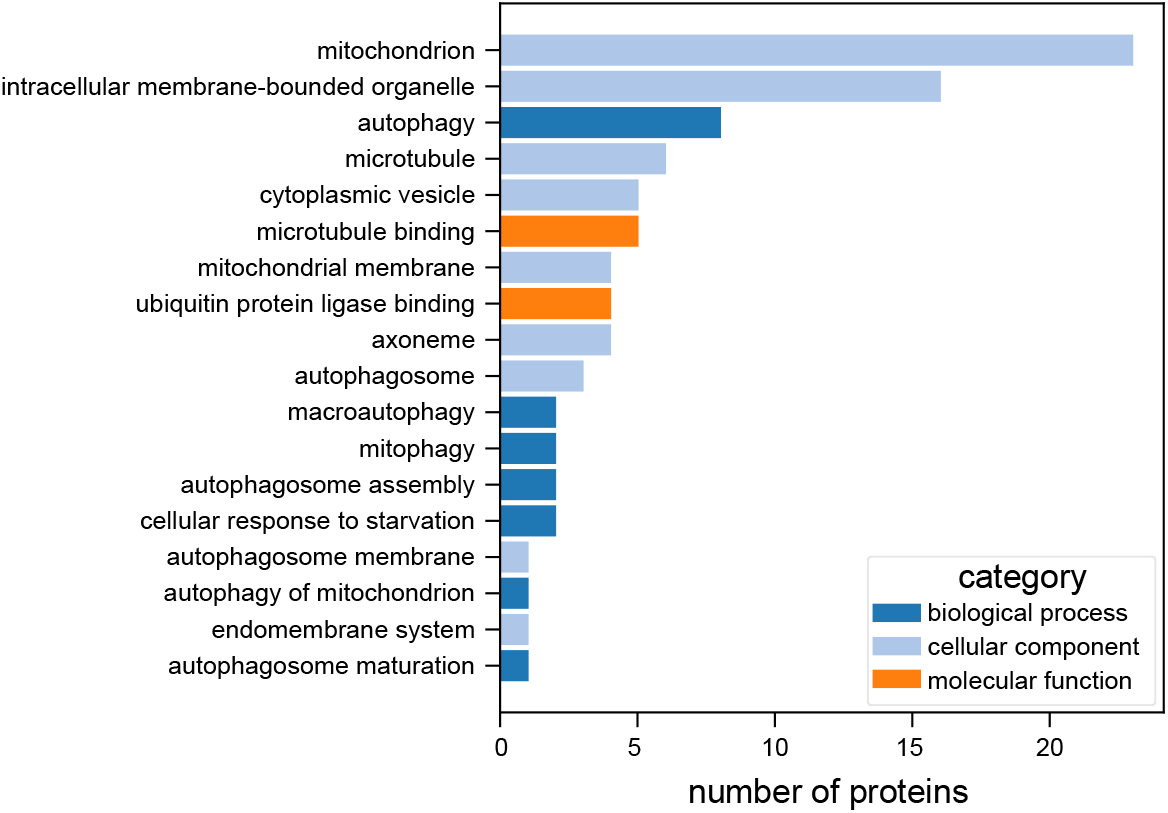
Gene Ontology (GO) terms shared between peptides in the HC-set and LC3B. GO terms shared between proteins with peptides in the HC-set and LC3B, colored by GO ontology category. Shared GO terms were sourced and processed from UniProt (The UniProt Consortium et al., 2025) (see Methods).

**Supplementary Figure 5.**
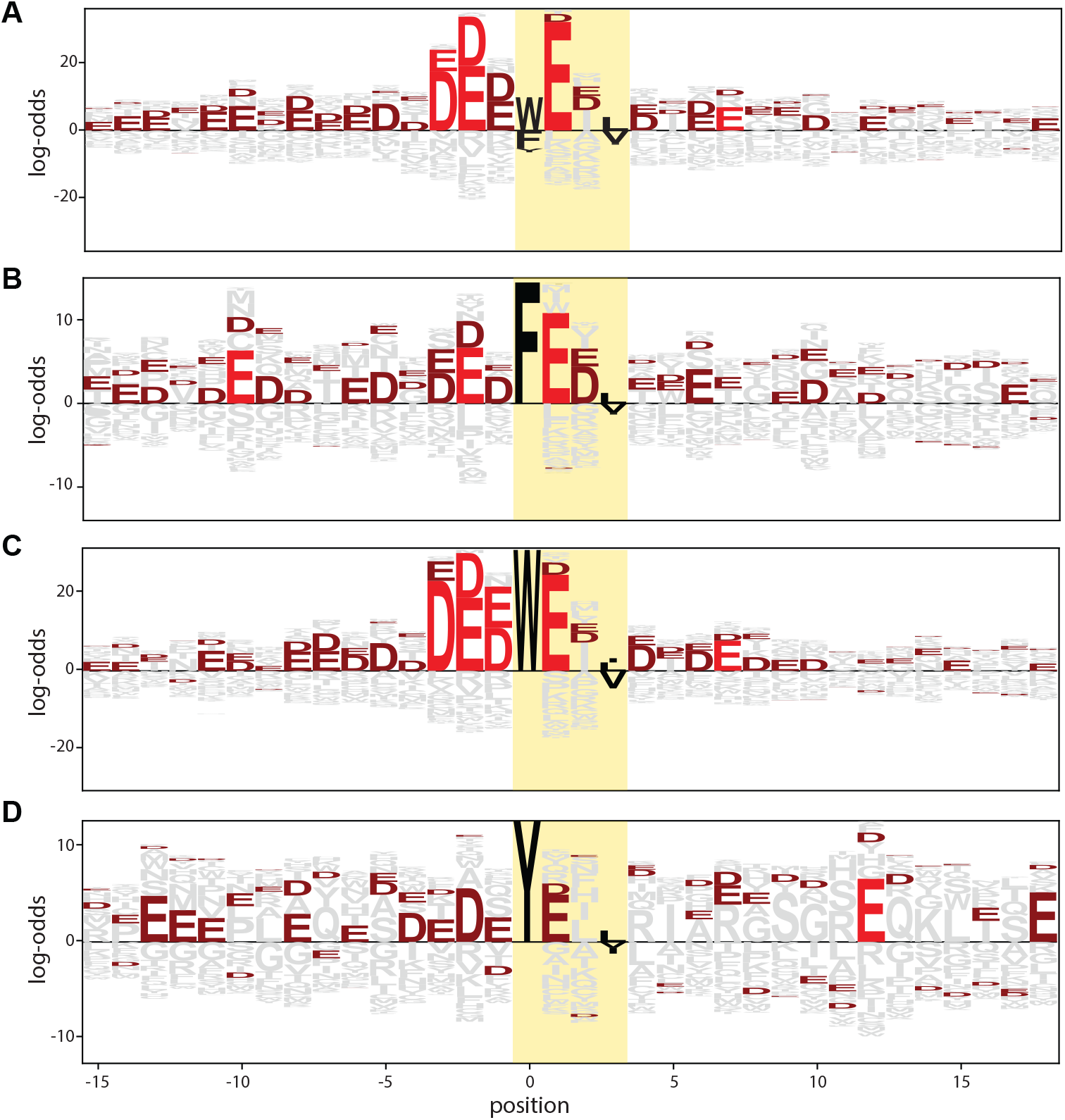
Sequence logos generated using hits with F-type, W-type, and Y-type LIRs. Sequence logos showing the enrichment or depletion of residues in peptides from the HC-set **(A)** or subsets of the HC-set bearing an F **(B)**, W **(C)**, or Y **(D)** residue in the X_0_ position. Log-odds ratio was calculated relative to all LIR-containing peptides in the input library using pLogo (O’Shea *et al*., 2013), with acidic residues colored maroon, and residues with statistically significant enrichment as calculated by pLogo in red. Core LIR motif positions are highlighted in yellow, with [FWY]_0_ and [LVI]_3_ in black.

**Supplementary Figure 6.**
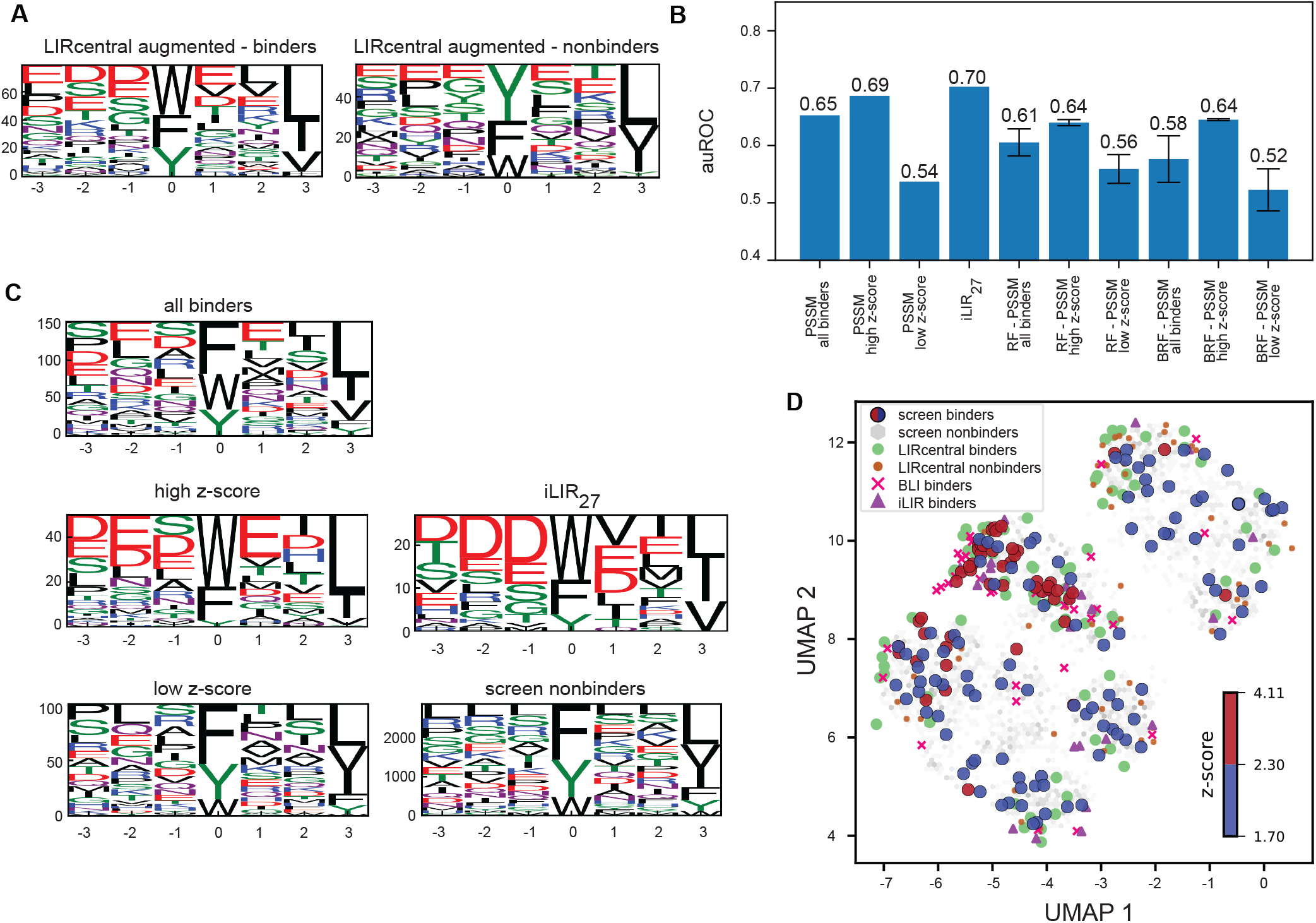
Sequence-based models to predict LC3B-binding peptides. **(A)** Sequence logos based on the LIRcentral augmented test set (see Methods). The height of each letter indicates the count of each residue at each position in the sequence set. **(B)** Performance of PSSM and Random Forest models in discriminating binding and non-binding sequences in the LIRcentral augmented test set. Performance was quantified by the area under the receiver operating characteristic (auROC) curve. Where present, error bars indicate standard deviation across randomly sampled subsets (see Methods). **(C)** Sequence logos for the screen-derived sequence sets and the 27 iLIR sequences used for model building and testing, as described in the methods (plotted as in A). **(D)** A UMAP plot (McInnes *et al*. 2018) of LIR sequence space, with each point marking a distinct sequence. Peptides from the HC-set are plotted as circles, colored based on z-score range. Peptides from the screen defined as non-binding are marked in light grey circles, and those defined as non-binding in LIRcentral are marked with orange circles. High z-score peptides from the screen (*i*.*e*., z-score ≥ 2.3; red circles) cluster with most of the LIRcentral (green) and iLIR27 (purple triangle) binders, whereas the more moderately enriching sequences (*i*.*e*., 1.7 ≤ z-score < 2.3; blue circles) occupy a much more diverse sequence space. Peptides measured to bind LC3B by BLI in this work are marked with a pink x.

**Supplementary Figure 7.**
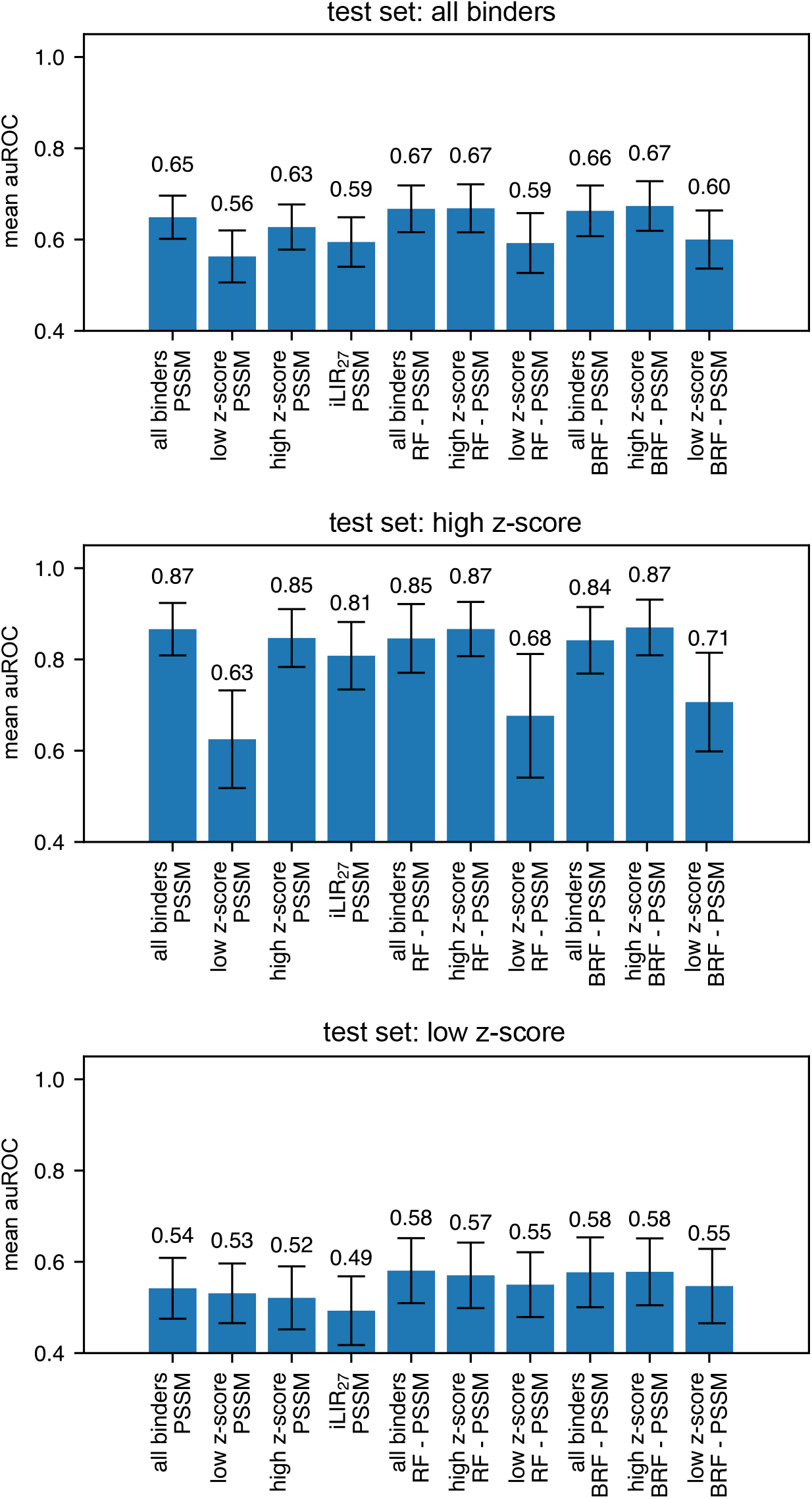
Model performance on test sets derived from screening data. Test sets were created from binder sequences sampled from different z-score ranges - top: all binder sequences (z-score ≥ 1.7); middle: high z-score sequences (z-score ≥ 2.3); bottom: low z-score sequences (1.7 ≤ z-score < 2.3) (see Methods). The y-axis is the mean auROC value across 100 replicate train/test splits, with error bars indicating the standard deviation.

**Supplementary Figure 8.**
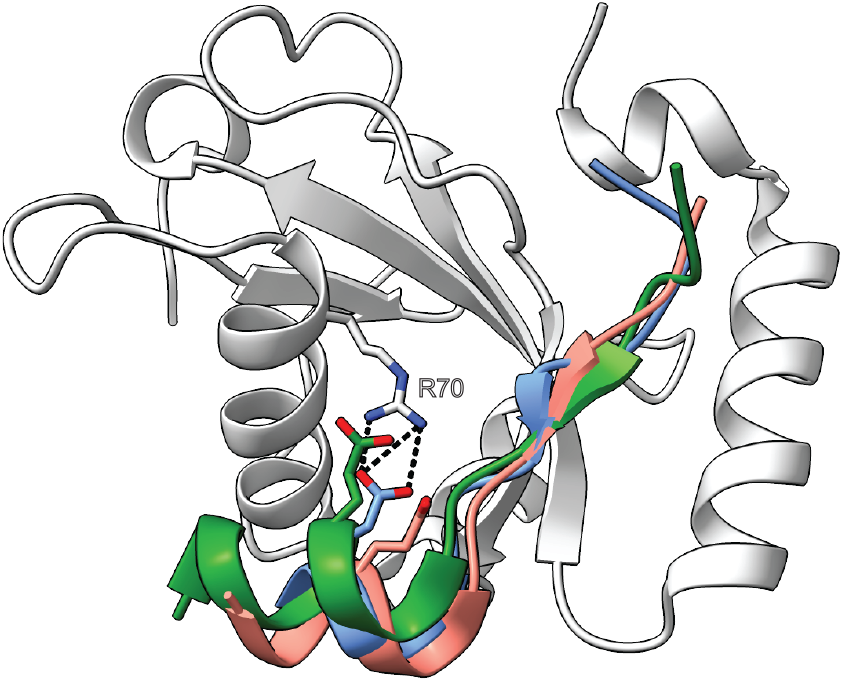
A glutamate in LC3B-binding peptides commonly contacts LC3B R70. Cartoon depiction of LC3B in complex with peptides from FYCO1^1276-1288^ (blue), ANK2^1588-1613^ (salmon), and ANK3^1985-2010^ (green), with glutamic acid residues from each peptide and LC3B residue R70 shown in stick representation. Contacts between FYCO1 and LC3B are shown in black dashes. The figure was generated by aligning LC3B from PDB entries 5d94 (Olsvik *et al*. 2015), 5yiq (Li *et al*. 2018), and 5cx3 (Cheng *et al*. 2016) to LC3B in 5d94 using ChimeraX (Meng *et al*. 2023), with only the LC3B of 5d94 shown for visual clarity.

**Supplementary Figure 9.**
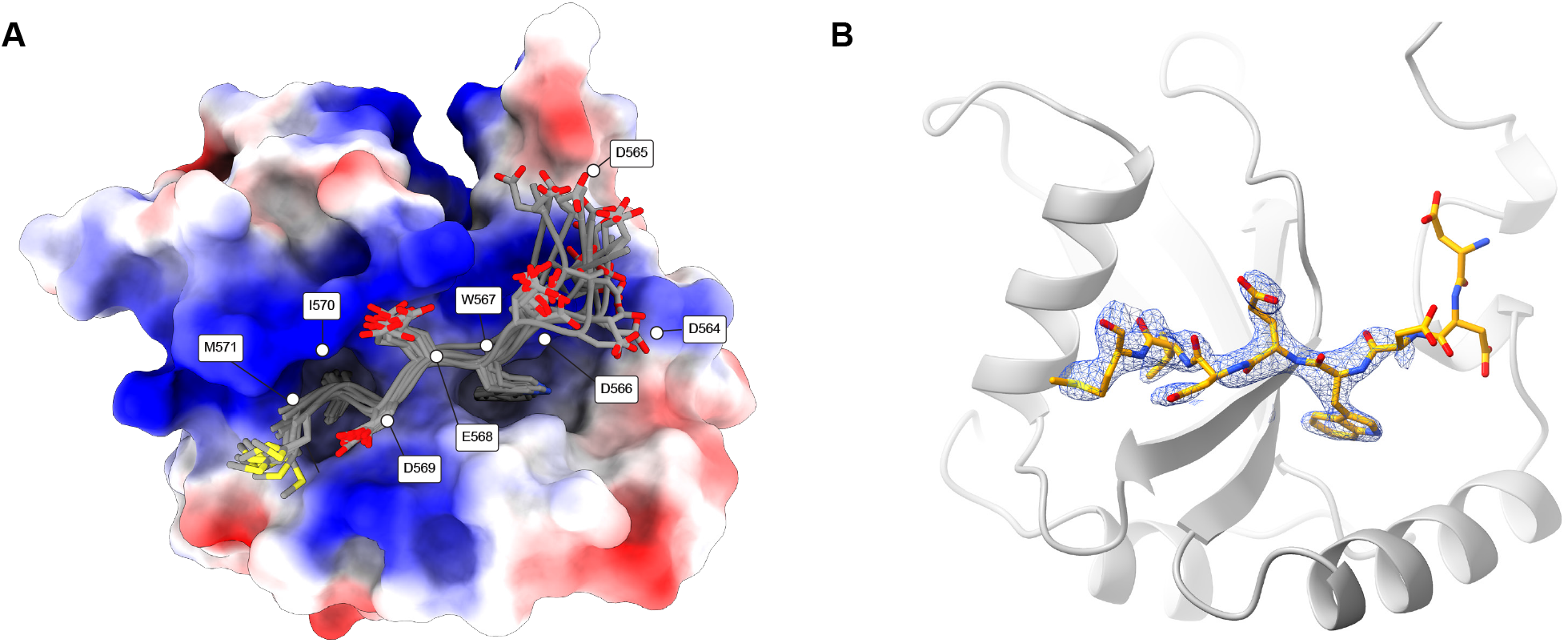
Ensemble refinement of the BLM-LC3B X-ray crystal structure. **(A)** Ensemble refinement (Burnley *et al*. 2012), using molecular dynamics refined against the X-ray data. The R_work_ and R_free_ for this ensemble are: 0.189 and 0.259; this R_free_ is lower than for the single-structure model (**Figure 3D-E, Supplementary Table 4**). Ten structures of the peptide (in stick representation) from the ensemble are shown, with LC3B using an electrostatic surface representation. The peptide N-terminus is less ordered than the LIR-core, consistent with the peptide interacting with the N-terminus of LC3B via multiple conformations. **(B)** Crystal structure of BLM^564-571^ (stick rendering) bound to LC3B (grey cartoon rendering) with 2m*Fo* − *DFc* electron density shown in blue mesh rendering, calculated with the BLM peptide atoms omitted during refinement, and contoured at 1σ.

**Supplementary Figure 10.**
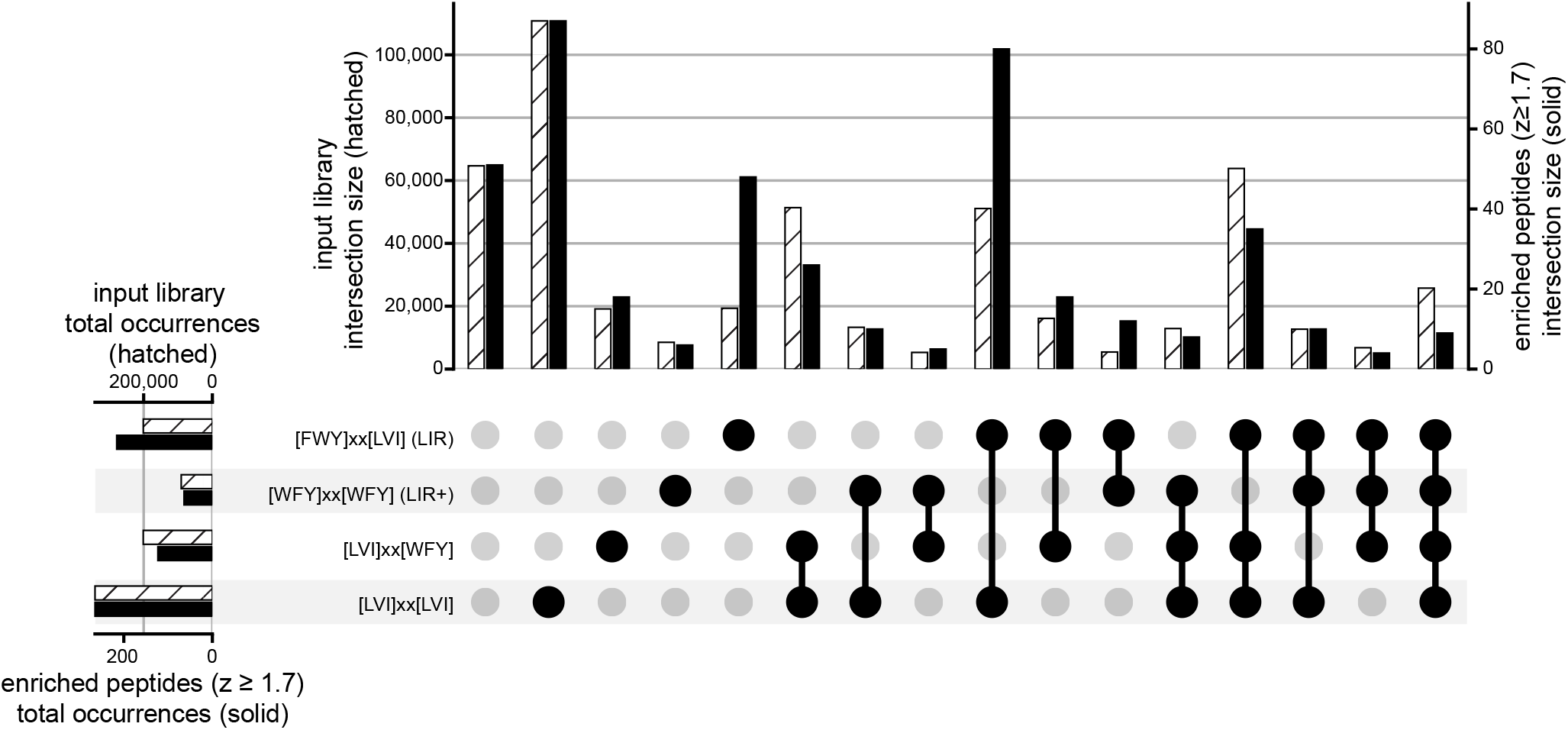
Presence of LIR and LIR-like motifs in the HC-set of peptides. Upset plot (Lex *et al*. 2014) with vertical bars marking the size of each intersection for the input library (hatched bars, left axis) and for enriched peptides with z ≥ 1.7 (solid bars, right axis). Canonical LIR motifs are defined as [FWY]xx[LVI] and LIR+ motifs as [WFY]xx[WFY], with x allowing for any amino acid. Additional motifs, [LVI]xx[LVI] and [LVI]xx[FWY], were included as AlphaFold3 (Abramson *et al*. 2024) predictions indicated the possibility of such motifs engaging LC3B. Peptides containing the biotin mimic “HPQ” motif (Devlin *et al*. 1990; Weber *et al*. 1992) were excluded from this analysis. Horizontal bars indicate the total number of occurrences of each motif in the input (hatched) and enriched (solid) peptide libraries.

**Supplementary Figure 11.**
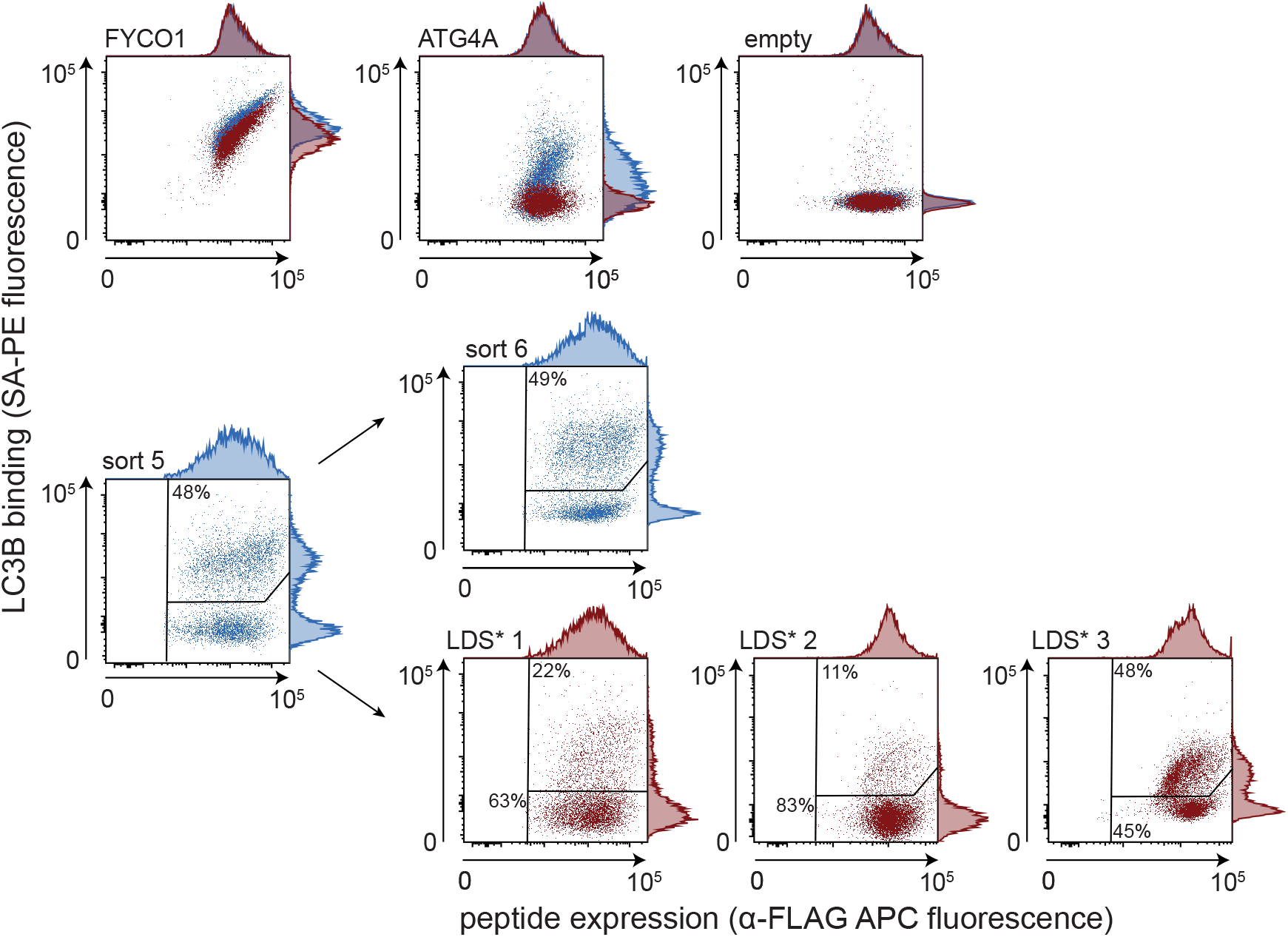
Bacterial display-based enrichment of peptides that bind to a pre-tetramerized LC3B LDS* mutant. An additional three rounds of bacterial display sorting were performed for binding to pre-tetramerized LC3B LDS*, using the population collected from round 5 of the LC3B sorting experiment as input. Signals for positive controls FYCO1^1264-1299^ and ATG4A^363-398^ and the empty-vector control are shown for 1.68 µM LC3B (blue) overlaid with 1.68 µM LC3B LDS* (red). The binding (upper right gate) and non-binding (lower right gate) populations for each LC3B LDS* sort, and a sixth round of LC3B binders, were collected and sequenced. The percentage of cells collected is indicated in each gate.

## Notes

### Competing Interest Statement

The authors have declared no competing interest.

### Summary of Updates

This version has been revised in response to peer review. We clarified portions of the text for accuracy and readability, including the Abstract, Results and Discussion sections. We added a supplementary figure panel showing the electron density omit map for the BLM peptide in our crystal structure. We also corrected minor text and reference-list errors. No new experimental data or conclusions were added.

## REFERENCES

Abramson J, Adler J, Dunger J, Evans R, Green T, Pritzel A, Ronneberger O, Willmore L, Ballard AJ, Bambrick J, et al. 2024. Accurate structure prediction of biomolecular interactions with AlphaFold 3. Nature 630: 493–500.

Abu-Jamous B, Kelly S. 2018. Clust: automatic extraction of optimal co-expressed gene clusters from gene expression data. Genome Biol 19: 172.

Behrends C, Sowa ME, Gygi SP, Harper JW. 2010. Network organization of the human autophagy system. Nature 466: 68–76.

Birgisdottir ÅB, Lamark T, Johansen T. 2013. The LIR motif - crucial for selective autophagy. J Cell Sci 126: 3237–3247.

Boecker CA, Holzbaur ELF. 2021. Hyperactive LRRK2 kinase impairs the trafficking of axonal autophagosomes. Autophagy 17: 2043–2045.

Brennan A, Layfield R, Long J, Williams HEL, Oldham NJ, Scott D, Searle MS. 2022. An ALS-associated variant of the autophagy receptor SQSTM1/p62 reprograms binding selectivity toward the autophagy-related hATG8 proteins. J Biol Chem 298: 101514.

Bugge K, Brakti I, Fernandes CB, Dreier JE, Lundsgaard JE, Olsen JG, Skriver K, Kragelund BB. 2020. Interactions by Disorder – A Matter of Context. Front Mol Biosci 7: 110.

Burnley BT, Afonine PV, Adams PD, Gros P. 2012. Modelling dynamics in protein crystal structures by ensemble refinement. eLife 1: e00311.

Bushnell B, Rood J, Singer E. 2017. BBMerge - Accurate paired shotgun read merging via overlap. PLoS One 12: e0185056.

Camacho C, Coulouris G, Avagyan V, Ma N, Papadopoulos J, Bealer K, Madden TL. 2009. BLAST+: architecture and applications. BMC Bioinformatics 10: 421.

Casadaban MJ, Cohen SN. 1980. Analysis of gene control signals by DNA fusion and cloning in Escherichia coli. Journal of Molecular Biology 138: 179–207.

Chatzichristofi A, Sagris V, Pallaris A, Eftychiou M, Kalvari I, Price N, Theodosiou T, Iliopoulos I, Nezis IP, Promponas VJ. 2023. LIRcentral: a manually curated online database of experimentally validated functional LIR motifs. Autophagy 19: 3189–3200.

Cheng X, Wang Y, Gong Y, Li F, Guo Y, Hu S, Liu J, Pan L. 2016. Structural basis of FYCO1 and MAP1LC3A interaction reveals a novel binding mode for Atg8-family proteins. Autophagy 12: 1330–1339.

Chiu JKH, Ong RT-H. 2022. Clustering biological sequences with dynamic sequence similarity threshold. BMC Bioinformatics 23: 108.

Cock PJA, Antao T, Chang JT, Chapman BA, Cox CJ, Dalke A, Friedberg I, Hamelryck T, Kauff F, Wilczynski B, et al. 2009. Biopython: freely available Python tools for computational molecular biology and bioinformatics. Bioinformatics 25: 1422–1423.

Davey NE, Simonetti L, Ivarsson Y. 2023. The next wave of interactomics: Mapping the SLiM-based interactions of the intrinsically disordered proteome. Current Opinion in Structural Biology 80: 102593.

Davey NE, Van Roey K, Weatheritt RJ, Toedt G, Uyar B, Altenberg B, Budd A, Diella F, Dinkel H, Gibson TJ. 2012. Attributes of short linear motifs. Mol BioSyst 8: 268–281.

Devlin JJ, Panganiban LC, Devlin PE. 1990. Random Peptide Libraries: a Source of Specific Protein Binding Molecules. Science 249: 404–406.

Diella F. 2008. Understanding eukaryotic linear motifs and their role in cell signaling and regulation. Front Biosci 13: 6580–6603.

DiMaio F, Echols N, Headd JJ, Terwilliger TC, Adams PD, Baker D. 2013. Improved low-resolution crystallographic refinement with Phenix and Rosetta. Nat Methods 10: 1102–1104.

Dyson HJ, Wright PE. 2005. Intrinsically unstructured proteins and their functions. Nat Rev Mol Cell Biol 6: 197–208.

Emsley P, Lohkamp B, Scott WG, Cowtan K. 2010. Features and development of Coot. Acta Crystallogr D Biol Crystallogr 66: 486–501.

Fas BA, Maiani E, Sora V, Kumar M, Mashkoor M, Lambrughi M, Tiberti M, Papaleo E. 2021. The conformational and mutational landscape of the ubiquitin-like marker for autophagosome formation in cancer. Autophagy 17: 2818–2841.

Florey O, Overholtzer M. 2012. Autophagy proteins in macroendocytic engulfment. Trends Cell Biol 22: 374–380.

Fumagalli F, Noack J, Bergmann TJ, Cebollero E, Pisoni GB, Fasana E, Fregno I, Galli C, Loi M, Soldà T, et al. 2016. Translocon component Sec62 acts in endoplasmic reticulum turnover during stress recovery. Nat Cell Biol 18: 1173–1184.

Genau HM, Huber J, Baschieri F, Akutsu M, Dötsch V, Farhan H, Rogov V, Behrends C. 2015. CUL3-KBTBD6/KBTBD7 Ubiquitin Ligase Cooperates with GABARAP Proteins to Spatially Restrict TIAM1-RAC1 Signaling. Molecular Cell 57: 995–1010.

Gibson DG, Young L, Chuang RY, Venter JC, Hutchison CA, Smith HO. 2009. Enzymatic assembly of DNA molecules up to several hundred kilobases. Nature Methods 6: 343–345.

Grant CE, Bailey TL. 2021. XSTREME: Comprehensive motif analysis of biological sequence datasets. bioRxiv. 2021.09.02.458722.

Gubas A, Dikic I. 2022. A guide to the regulation of selective autophagy receptors. FEBS J 289: 75–89.

Heckmann BL, Teubner BJW, Tummers B, Boada-Romero E, Harris L, Yang M, Guy CS, Zakharenko SS, Green DR. 2019. LC3-Associated Endocytosis Facilitates β-Amyloid Clearance and Mitigates Neurodegeneration in Murine Alzheimer’s Disease. Cell 178: 536–551.e14.

Hwang T, Parker SS, Hill SM, Grant RA, Ilunga MW, Sivaraman V, Mouneimne G, Keating AE. 2022. Native proline-rich motifs exploit sequence context to target actin-remodeling Ena/VASP protein ENAH. eLife 11: e70680.

Johansen T, Lamark T. 2020. Selective Autophagy: ATG8 Family Proteins, LIR Motifs and Cargo Receptors. J Mol Biol 432: 80–103.

Jumper J, Evans R, Pritzel A, Green T, Figurnov M, Ronneberger O, Tunyasuvunakool K, Bates R, Žídek A, Potapenko A, et al. 2021. Highly accurate protein structure prediction with AlphaFold. Nature 596: 583–589.

Kabeya Y, Mizushima N, Yamamoto A, Oshitani-Okamoto S, Ohsumi Y, Yoshimori T. 2004. LC3, GABARAP and GATE16 localize to autophagosomal membrane depending on form-II formation. J Cell Sci 117: 2805–2812.

Kabsch W. 2010. XDS. Acta Crystallogr D Biol Crystallogr 66: 125–132.

Kalvari I, Tsompanis S, Mulakkal NC, Osgood R, Johansen T, Nezis IP, Promponas VJ. 2014. iLIR: A web resource for prediction of Atg8-family interacting proteins. Autophagy 10: 913–925.

Katz BA. 1995. Binding to protein targets of peptidic leads discovered by phage display: crystal structures of streptavidin-bound linear and cyclic peptide ligands containing the HPQ sequence. Biochemistry 34: 15421–15429.

Kliche J, Garvanska DH, Simonetti L, Badgujar D, Dobritzsch D, Nilsson J, Davey NE, Ivarsson Y. 2023. Large-scale phosphomimetic screening identifies phospho-modulated motif-based protein interactions. Mol Syst Biol 19: e11164.

Kliche J, Ivarsson Y. 2022. Orchestrating serine/threonine phosphorylation and elucidating downstream effects by short linear motifs. Biochem J 479: 1–22.

Knævelsrud H, Carlsson SR, Simonsen A. 2013. SNX18 tubulates recycling endosomes for autophagosome biogenesis. Autophagy 9: 1639–1641.

Kraft LJ, Nguyen TA, Vogel SS, Kenworthy AK. 2014. Size, stoichiometry, and organization of soluble LC3-associated complexes. Autophagy 10: 861–877.

Kumar M, Michael S, Alvarado-Valverde J, Mészáros B, Sámano-Sánchez H, Zeke A, Dobson L, Lazar T, Örd M, Nagpal A, et al. 2022. The Eukaryotic Linear Motif resource: 2022 release. Nucleic Acids Research 50: D497–D508.

Kumar M, Michael S, Alvarado-Valverde J, Zeke A, Lazar T, Glavina J, Nagy-Kanta E, Donagh JM, Kalman ZE, Pascarelli S, et al. 2024. ELM—the Eukaryotic Linear Motif resource—2024 update. Nucleic Acids Research 52: D442–D455.

Larman HB, Zhao Z, Laserson U, Li MZ, Ciccia A, Gakidis MAM, Church GM, Kesari S, LeProust EM, Solimini NL, et al. 2011. Autoantigen discovery with a synthetic human peptidome. Nat Biotechnol 29: 535–541.

Lee A, Davis JH. 2024. NCOA4 initiates ferritinophagy by binding GATE16 using two highly avid short linear interaction motifs. bioRxiv 2024.06.09.597909.

Lemaître G, Nogueira F, Aridas CK. 2017. Imbalanced-learn: a python toolbox to tackle the curse of imbalanced datasets in machine learning. J Mach Learn Res 18: 559–563.

Lex A, Gehlenborg N, Strobelt H, Vuillemot R, Pfister H. 2014. UpSet: Visualization of Intersecting Sets. IEEE Trans Vis Comput Graph 20: 1983–1992.

Li J, Zhu R, Chen K, Zheng H, Zhao H, Yuan C, Zhang H, Wang C, Zhang M. 2018. Potent and specific Atg8-targeting autophagy inhibitory peptides from giant ankyrins. Nat Chem Biol 14: 778–787.

Liebschner D, Afonine PV, Baker ML, Bunkóczi G, Chen VB, Croll TI, Hintze B, Hung L-W, Jain S, McCoy AJ, et al. 2019. Macromolecular structure determination using X-rays, neutrons and electrons: recent developments in Phenix. Acta Crystallogr D Struct Biol 75: 861–877.

Liu H, Naismith JH. 2008. An efficient one-step site-directed deletion, insertion, single and multiple-site plasmid mutagenesis protocol. BMC Biotechnology 8: 91.

Lv M, Wang C, Li F, Peng J, Wen B, Gong Q, Shi Y, Tang Y. 2017. Structural insights into the recognition of phosphorylated FUNDC1 by LC3B in mitophagy. Protein & Cell 8: 25–38.

Madureira M, Connor-Robson N, Wade-Martins R. 2020. LRRK2: Autophagy and Lysosomal Activity. Front Neurosci 14.

Marshall RS, Hua Z, Mali S, McLoughlin F, Vierstra RD. 2019. ATG8-Binding UIM Proteins Define a New Class of Autophagy Adaptors and Receptors. Cell 177: 766–781.e24.

McCoy AJ, Grosse-Kunstleve RW, Adams PD, Winn MD, Storoni LC, Read RJ. 2007. Phaser crystallographic software. J Appl Crystallogr 40: 658–674.

McInnes, L., Healy, J. and Melville, J. 2018. UMAP: Uniform Manifold Approximation and Projection for Dimension Reduction. arXiv.

Meng EC, Goddard TD, Pettersen EF, Couch GS, Pearson ZJ, Morris JH, Ferrin TE. 2023. UCSF ChimeraX: Tools for structure building and analysis. Protein Science 32: e4792.

Mizushima N, Levine B, Cuervo AM, Klionsky DJ. 2008. Autophagy fights disease through cellular self-digestion. Nature 451: 1069–1075.

Montes-Fernández MA, Pérez-Villegas EM, Garcia-Gonzalo FR, Pedrazza L, Rosa JL, de Toledo GA, Armengol JA. 2020. The HERC1 ubiquitin ligase regulates presynaptic membrane dynamics of central synapses. Sci Rep 10: 12057.

Olsvik HL, Lamark T, Takagi K, Larsen KB, Evjen G, Øvervatn A, Mizushima T, Johansen T. 2015. FYCO1 Contains a C-terminally Extended, LC3A/B-preferring LC3-interacting Region (LIR) Motif Required for Efficient Maturation of Autophagosomes during Basal Autophagy. J Biol Chem 290: 29361–29374.

O’Shea JP, Chou MF, Quader SA, Ryan JK, Church GM, Schwartz D. 2013. pLogo: a probabilistic approach to visualizing sequence motifs. Nat Methods 10: 1211–1212.

Pankiv S, Clausen TH, Lamark T, Brech A, Bruun J-A, Outzen H, Øvervatn A, Bjørkøy G, Johansen T. 2007. p62/SQSTM1 binds directly to Atg8/LC3 to facilitate degradation of ubiquitinated protein aggregates by autophagy. J Biol Chem 282: 24131–24145.

Park H, Kang J-H, Lee S. 2020. Autophagy in Neurodegenerative Diseases: A Hunter for Aggregates. IJMS 21: 3369.

Park S, Han S, Choi I, Kim B, Park SP, Joe E-H, Suh YH. 2016. Interplay between Leucine-Rich Repeat Kinase 2 (LRRK2) and p62/SQSTM-1 in Selective Autophagy. PLoS One 11: e0163029.

Pedregosa F, Varoquaux G, Gramfort A, Michel V, Thirion B, Grisel O, Blondel M, Prettenhofer P, Weiss R, Dubourg V, et al. 2011. Scikit-learn: Machine Learning in Python. Journal of Machine Learning Research 12: 2825–2830.

Pérez-Villegas EM, Pérez-Rodríguez M, Negrete-Díaz JV, Ruiz R, Rosa JL, de Toledo GA, Rodríguez-Moreno A, Armengol JA. 2020. HERC1 Ubiquitin Ligase Is Required for Hippocampal Learning and Memory. Front Neuroanat 14: 592797.

Popelka H. 2020. Dancing while self-eating: Protein intrinsic disorder in autophagy. Prog Mol Biol Transl Sci 174: 263–305.

Qiu Y, Zheng Y, Wu K-P, Schulman BA. 2017. Insights into links between autophagy and the ubiquitin system from the structure of LC3B bound to the LIR motif from the E3 ligase NEDD4. Protein Sci 26: 1674–1680.

Ramesh Babu J, Lamar Seibenhener M, Peng J, Strom A, Kemppainen R, Cox N, Zhu H, Wooten MC, Diaz-Meco MT, Moscat J, et al. 2008. Genetic inactivation of p62 leads to accumulation of hyperphosphorylated tau and neurodegeneration. Journal of Neurochemistry 106: 107–120.

Rice JJ, Schohn A, Bessette PH, Boulware KT, Daugherty PS. 2006. Bacterial display using circularly permuted outer membrane protein OmpX yields high affinity peptide ligands. Protein Sci 15: 825–836.

Richter B, Sliter DA, Herhaus L, Stolz A, Wang C, Beli P, Zaffagnini G, Wild P, Martens S, Wagner SA, et al. 2016. Phosphorylation of OPTN by TBK1 enhances its binding to Ub chains and promotes selective autophagy of damaged mitochondria. Proc Natl Acad Sci U S A 113: 4039–4044.

Rogov VV, Suzuki H, Marinković M, Lang V, Kato R, Kawasaki M, Buljubašić M, Šprung M, Rogova N, Wakatsuki S, Hamacher-Brady A, Dötsch V, Dikic I, Brady NR, Novak I. 2017. Phosphorylation of the mitochondrial autophagy receptor Nix enhances its interaction with LC3 proteins. Sci Rep. 7(1):1131.

Rogov VV, Nezis IP, Tsapras P, Zhang H, Dagdas Y, Noda NN, Nakatogawa H, Wirth M, Mouilleron S, McEwan DG, et al. 2023. Atg8 family proteins, LIR/AIM motifs and other interaction modes. Autophagy Rep 2: 2188523.

Rogov VV, Suzuki H, Fiskin E, Wild P, Kniss A, Rozenknop A, Kato R, Kawasaki M, McEwan DG, Löhr F, et al. 2013. Structural basis for phosphorylation-triggered autophagic clearance of Salmonella. Biochemical Journal 454: 459–466.

Roosen DA, Cookson MR. 2016. LRRK2 at the interface of autophagosomes, endosomes and lysosomes. Mol Neurodegener 11: 73.

Rozenknop A, Rogov VV, Rogova NYu, Löhr F, Güntert P, Dikic I, Dötsch V. 2011. Characterization of the Interaction of GABARAPL-1 with the LIR Motif of NBR1. Journal of Molecular Biology 410: 477–487.

Rubin AF, Gelman H, Lucas N, Bajjalieh SM, Papenfuss AT, Speed TP, Fowler DM. 2017. A statistical framework for analyzing deep mutational scanning data. Genome Biol 18: 150.

Sawa-Makarska J, Abert C, Romanov J, Zens B, Ibiricu I, Martens S. 2014. Cargo binding to Atg19 unmasks additional Atg8 binding sites to mediate membrane-cargo apposition during selective autophagy. Nat Cell Biol 16: 425–433.

Schneider TD, Stephens RM. 1990. Sequence logos: a new way to display consensus sequences. Nucleic Acids Res 18: 6097–6100.

Skytte Rasmussen M, Mouilleron S, Kumar Shrestha B, Wirth M, Lee R, Bowitz Larsen K, Abudu Princely Y, O’Reilly N, Sjøttem E, Tooze SA, et al. 2017. ATG4B contains a C-terminal LIR motif important for binding and efficient cleavage of mammalian orthologs of yeast Atg8. Autophagy 13: 834–853.

Sønder SL, Häger SC, Heitmann ASB, Frankel LB, Dias C, Simonsen AC, Nylandsted J. 2021. Restructuring of the plasma membrane upon damage by LC3-associated macropinocytosis. Sci Adv 7: eabg1969.

Stark C, Breitkreutz B-J, Reguly T, Boucher L, Breitkreutz A, Tyers M. 2006. BioGRID: a gen al repository for interaction datasets. Nucleic Acids Res 34: D535–539.

Stadel D, Millarte V, Tillmann KD, Huber J, Tamin-Yecheskel B-C, Akutsu M, Demishtein A, Ben-Zeev B, Anikster Y, Perez F, et al. 2015. TECPR2 Cooperates with LC3C to Regulate COPII-Dependent ER Export. Molecular Cell 60: 89–104.

Sugawara K, Suzuki NN, Fujioka Y, Mizushima N, Ohsumi Y, Inagaki F. 2004. The crystal structure of microtubule-associated protein light chain 3, a mammalian homologue of Saccharomyces cerevisiae Atg8. Genes to Cells 9: 611–618.

Tareen A, Kinney JB. 2020. Logomaker: beautiful sequence logos in Python. Bioinformatics 36: 2272–2274.

The Gene Ontology Consortium, Aleksander SA, Balhoff J, Carbon S, Cherry JM, Drabkin HJ, Ebert D, Feuermann M, Gaudet P, Harris NL, et al. 2023. The Gene Ontology knowledgebase in 2023. Genetics 224: iyad031.

The UniProt Consortium, Bateman A, Martin M-J, Orchard S, Magrane M, Adesina A, Ahmad S, Bowler-Barnett EH, Bye-A-Jee H, Carpentier D, et al. 2025. UniProt: the Universal Protein Knowledgebase in 2025. Nucleic Acids Research 53: D609–D617.

Tompa P, Davey NE, Gibson TJ, Babu MM. 2014. A Million Peptide Motifs for the Molecular Biologist. Molecular Cell 55: 161–169.

Tong Y, Zhu W, Chen J, Zhang W, Xu F, Pang J. 2023. Targeted Degradation of Alpha-Synuclein by Autophagosome-Anchoring Chimera Peptides. J Med Chem 66: 12614–12628.

Tsao KL, DeBarbieri B, Michel H, Waugh DS. 1996. A versatile plasmid expression vector for the production of biotinylated proteins by site-specific, enzymatic modification in Escherichia coli. Gene 169: 59–64.

Van Roey K, Uyar B, Weatheritt RJ, Dinkel H, Seiler M, Budd A, Gibson TJ, Davey NE. 2014. Short Linear Motifs: Ubiquitous and Functionally Diverse Protein Interaction Modules Directing Cell Regulation. Chem Rev 114: 6733–6778.

Weber PC, Pantoliano MW, Thompson LD. 1992. Crystal structure and ligand-binding studies of a screened peptide complexed with streptavidin. Biochemistry 31: 9350–9354.

Wirth, M., Zhang, W., Razi, M. et al. 2019. Molecular determinants regulating selective binding of autophagy adapters and receptors to ATG8 proteins. Nat Commun 10, 2055.

Wirth M, Mouilleron S, Zhang W, Sjøttem E, Princely Abudu Y, Jain A, Lauritz Olsvik H, Bruun J-A, Razi M, Jefferies HBJ, et al. 2021. Phosphorylation of the LIR Domain of SCOC Modulates ATG8 Binding Affinity and Specificity. J Mol Biol 433: 166987.

Wright PE, Dyson HJ. 2015. Intrinsically disordered proteins in cellular signalling and regulation. Nat Rev Mol Cell Biol 16: 18–29.

Wurzer B, Zaffagnini G, Fracchiolla D, Turco E, Abert C, Romanov J, Martens S. 2015. Oligomerization of p62 allows for selection of ubiquitinated cargo and isolation membrane during selective autophagy. eLife 4: e08941.

Zaffagnini G, Martens S. 2016. Mechanisms of Selective Autophagy. J Mol Biol 428: 1714–1724.

